# Repeat modules and N-linked glycans define structure and antigenicity of a critical enterotoxigenic *E. coli adhesin*

**DOI:** 10.1101/2024.05.08.593125

**Authors:** Zachary T. Berndsen, Marjahan Akhtar, Mahima Thapa, Tim Vickers, Aaron Schmitz, Jonathan L. Torres, Sabyasachi Baboo, Pardeep Kumar, Nazia Khatoom, Alaullah Sheikh, Melissa Hamrick, Jolene K. Diedrich, Salvador Martinez-Bartolome, Patrick T. Garrett, John R. Yates, Jackson S. Turner, Renee M. Laird, Frédéric Poly, Chad K. Porter, Jeffrey Copps, Ali H. Ellebedy, Andrew B. Ward, James M. Fleckenstein

## Abstract

Enterotoxigenic *Escherichia coli* (ETEC) cause hundreds of millions of cases of infectious diarrhea annually, predominantly in children from low-middle income regions. Notably, in children, as well as human volunteers challenged with ETEC, diarrheal severity is significantly increased severity in blood group A (bgA) individuals. EtpA, is a secreted glycoprotein adhesin that functions as a blood group A lectin to promote critical interactions between ETEC and blood group A glycans on intestinal epithelia for effective bacterial adhesion and toxin delivery. EtpA is highly immunogenic resulting in robust antibody responses following natural infection and experimental challenge of human volunteers with ETEC. To understand how EtpA directs ETEC-blood group A interactions and stimulates adaptive immunity, we mutated EtpA, mapped its glycosylation by mass-spectrometry (MS), isolated polyclonal (pAbs) and monoclonal antibodies (mAbs) from vaccinated mice and ETEC-infected human volunteers, and determined structures of antibody-EtpA complexes by cryo-electron microscopy. Both bgA and mAbs that inhibited EtpA-bgA interactions and ETEC adhesion, bound to the C-terminal repeat domain highlighting this region as crucial for ETEC pathogen-host interaction. MS analysis uncovered extensive and heterogeneous N-linked glycosylation of EtpA and cryo-EM structures revealed that mAbs directly engage these unique glycan containing epitopes. Finally, electron microscopy-based polyclonal epitope mapping revealed antibodies targeting numerous distinct epitopes on N and C-terminal domains, suggesting that EtpA vaccination generates responses against neutralizing and decoy regions of the molecule. Collectively, we anticipate that these data will inform our general understanding of pathogen-host glycan interactions and adaptive immunity relevant to rational vaccine subunit design.

**Author summary:** Enterotoxigenic *E. coli* (ETEC), a leading cause of diarrhea disproportionately affecting young children in low-income regions, are a priority for vaccine development. Individuals possessing A blood-type are more susceptible to severe cholera-like disease. EtpA, a secreted, immunogenic, blood group A binding protein, is a current vaccine target antigen. Here, we determined the atomic structure of EtpA in complex with protective as well as non-protective monoclonal antibodies targeting two different domains of the protein, allowing us to pinpoint key regions involved in blood-group A antigen recognition and uncover the mechanism of antibody-based protection. In addition, we show through mass-spectrometry that EtpA is extensively and heterogeneously glycosylated at surface-exposed asparagine residues by a promiscuous and low-fidelity glycosyltransferase, EtpC, and that this unique form of bacterial glycosylation is critical for to development of protective immune responses. Lastly, polyclonal antibodies from vaccinated mice as well as monoclonal antibodies obtained from ETEC-infected human volunteers revealed that the highly antigenic surface of EtpA exhibits both protective and non-protective epitopes. These results greatly expand our understanding of ETEC pathogenesis, and the immune responses elicited by these common infections, providing valuable information to aid in the rational design and testing of subunit vaccines.

## Introduction

Enterotoxigenic *Escherichia coli* (ETEC) are diarrheal pathogens defined by their production of heat-labile (LT) and heat-stable (ST) enterotoxins^1^. ETEC, an exceedingly common cause of infectious diarrhea in areas where clean water and sanitation remain limited, accounts for hundreds of millions of cases of acute diarrheal illness each year^2^. In addition, these pathogens are a leading cause of more severe diarrhea and death^3,4^ among young children of low-income regions and are associated with long-term sequelae including poor growth^5–9^ and malnutrition^10–13^.

Given the persistent and pervasive impacts of ETEC infections, these pathogens have remained a high priority for vaccine development^14–16^. Efforts to identify novel surface-expressed molecules that might be targeted in ETEC vaccine development led to the identification of the plasmid-borne *etpBAC* two-partner secretion (TPS) locus responsible for export of EtpA, an extracellular adhesin^17^. The *etpBAC* locus encodes EtpB a polypeptide-transport-associated (POTRA)^18^ domain (TpsB) transmembrane protein required for EtpA secretion, the extracellular EtpA (TpsA) adhesin, and EtpC, a glycosyltransferase responsible for glycosylation of EtpA^17^. All three genes are required for optimal secretion of EtpA. The EtpA molecule is typically heavily glycosylated and *etpC* mutants exhibit dramatically reduced production of EtpA, as well as altered tropism for target epithelial cells, suggesting that glycosylation of EtpA may be important for proper folding and function of the adhesin^17^.

Once secreted, the high molecular weight (~170 kDa) EtpA glycoprotein serves as a unique molecular bridge between the bacteria and intestinal mucosal surfaces^19^, essential to pathogen-host interactions required for delivery of both LT^20,21^ and ST^22^. On host epithelia, EtpA binds to N-acetylgalactosamine (GalNAc) residues on enterocyte surfaces as well as secreted mucins including MUC2, interactions that are critical for efficient adhesion, toxin delivery, and intestinal colonization^23^. EtpA preferentially engages GalNAc as the terminal sugar of human A blood group presented on enterocytes. Importantly, human volunteers challenged with the EtpA-producing H10407 strain of ETEC were significantly more likely to develop moderate-severe diarrhea if they were blood group A^24^, recapitulating earlier observations that young bgA+ children in Bangladesh were more likely to develop diarrhea with ETEC infection^9^.

In exploring the utility of EtpA as a potential vaccine antigen, studies to date have demonstrated that the *etpBAC* locus is highly conserved across ETEC from geographically disparate origins^25–30^, is immunogenic following natural^27,30^ and human experimental challenge^31,32^ infections, and that immunization with recombinant EtpA (rEtpA) affords considerable protection against intestinal colonization^21,27,29,33,19,34–37^. In addition, EtpA expression by ETEC strains is strongly associated with the development of diarrheal illness in young children, while antibodies against EtpA are associated with protection^30^. Despite enthusiasm for targeting EtpA in next-generation ETEC vaccines,^38–40^ relatively little is known about its structure, antigenicity, or the mechanisms by which antibodies targeting this molecule mediate protection.

Several structures of truncated N-terminal (TPS) domains required for secretion of TpsA molecules ^41–45^ as well as a single full-length TpsA exoprotein^46^, have been solved by X-ray crystallography. The highly homologous TPS domain structures all adopt a similar fold, specifically, an extended 3-sided β-helix. These N-terminal TPS domains may be followed by a series of repeat modules, with some such as HxuA^46^ containing short extra-helical loops or motifs thought to contribute to function ^47^. The N-terminal secretion domain of EtpA was previously shown to be sufficient for export as well as binding to flagella^48^, while the function of a series of four C-terminal repeats was unknown.

Here we provide the complete structure of EtpA determined by cryo-EM, both alone and complexed to anti-EtpA monoclonal antibodies produced by vaccination, as well as high-resolution mass-spectrometry (MS) analysis of EtpA glycosylation. These data define regions of the molecule that are required for activity and are targets for antibody neutralization, provide a detailed profile of its extensive antigenic glycosylation, and show that alterations in glycosylation can impact antibody function. We identified a diverse set of epitopes from polyclonal sera of vaccinated mice, indicating that EtpA possesses a large and variable immunogenic surface with numerous potential neutralizing and decoy epitopes. We anticipate that the elucidation of the antigenic structure of this important virulence factor will afford insights into molecular correlates of protection and help guide further development of EtpA and similar proteins as vaccine immunogens.

## Results

### EtpA repeat regions direct blood group A binding on target host cells

Like many bacterial adhesins, EtpA is a lectin, or a carbohydrate binding protein^23^. Similar to another TpsA protein, filamentous hemagglutinin of *Bordetella pertussis*^49,50^, EtpA also possesses the ability to agglutinate erythrocytes. Hemagglutination activity of lectins typically arises when these molecules possess two or more carbohydrate binding sites permitting cross-linking of cells^51^. While individual interactions may be of low affinity, high avidity can be achieved through tandem repetition of lectin-binding regions^52–54^. To determine whether the C-terminal region of EtpA, comprised of 4 repeat modules (figure 1A), was involved in blood group A glycan recognition, we first examined a truncated version of recombinant EtpA (rEtpA_1-1086_, figure 1A), lacking the full complement of repeat modules. We found that while the truncated molecule was efficiently secreted, it was incapable of binding efficiently to blood group A glycans on the surface of intestinal epithelial cells (figure 1B), on solid substrates (figure 1C), or erythrocytes (figure 1D) suggesting that the repeats act in concert to engage target carbohydrates.

**Figure 1.**
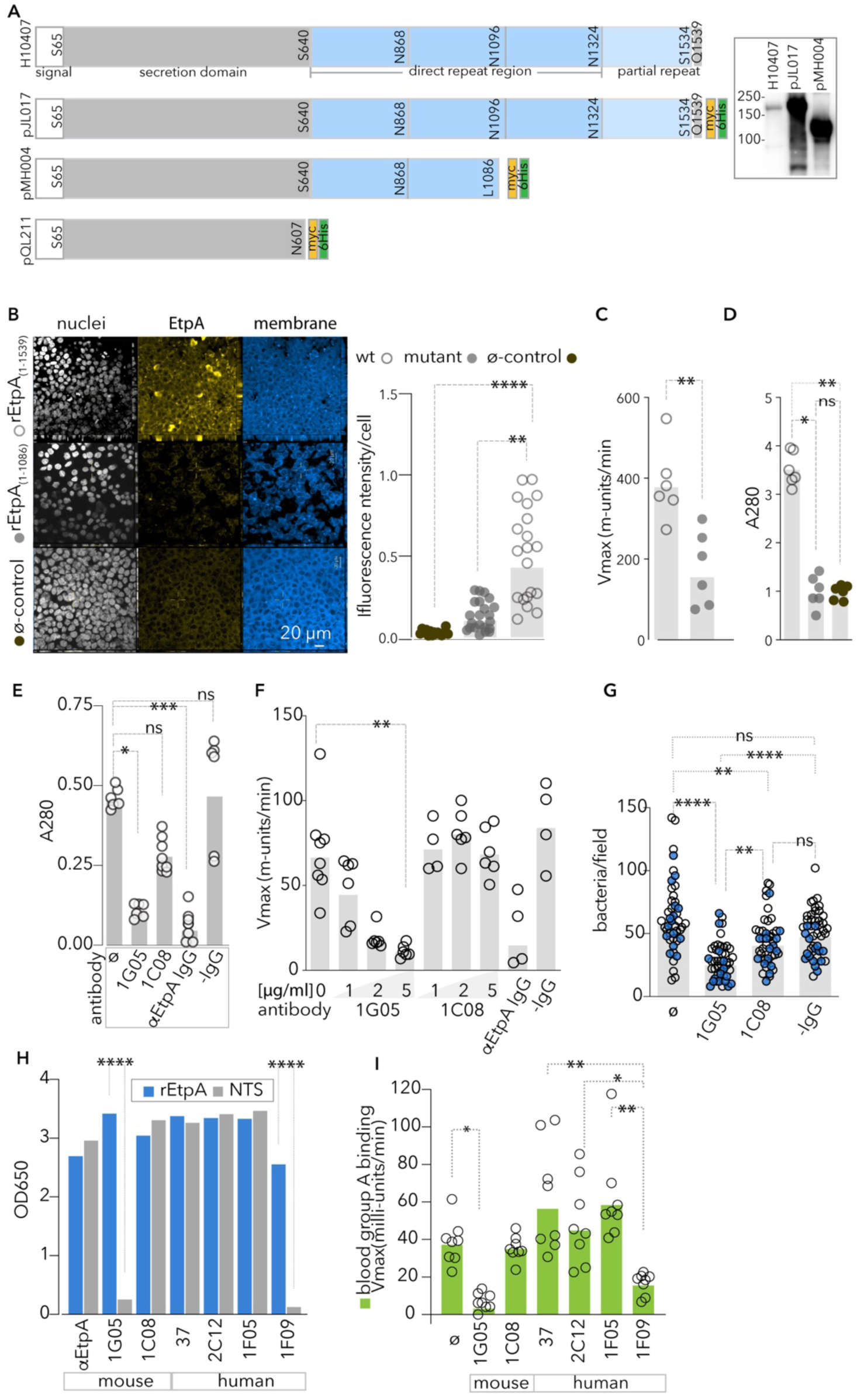
The repeat region of EtpA directs critical interactions with A blood group glycans. **A.** Schematic depicts molecular organization of the EtpA molecule from ETEC H10407 (top), recombinant EtpA encoded on pJY017, and the truncated recombinant antigens EtpA (1-1086) and the NTS domain (1-607) encoded on pMH004, and pQL211, respectively. Inset: anti-EtpA immunoblot of TCA-precipitated culture supernatants from H10407 wild type strain, and recombinant Top10 strains jf1696 and jf5090 carrying plasmids pJL017 and pMH004, respectively. **B.** Binding of full-length and truncated, mutant EtpA molecules to blood group A-expressing HT-29 cells. Shown at left are representative fields quantified in graph at right from n-20 replicate fields from two independent experiments. Bars indicate geometric mean fluorescence intensity per cell (****≤0.0001, **=0.0015 by Kruskal-Wallis nonparametric testing). **C.** Kinetic ELISA data reflect binding of full-length and truncated EtpA to blood group A **=0.0043 (Mann-Whitney, two-tailed). **D.** Blood group A1 erythrocyte (A1-RBC) pull-down assay with full-length and truncated EtpA. *=0.0147, **0.0073 by Kruskal-Wallis. Results in C,D represent combination of technical duplicates from three independent experiments. **E**. Inhibition of EtpA-A1-RBC interactions with anti-EtpA monoclonal antibodies recognizing the repeat (mAb 1G05) and secretion (mAb 1G08) domains compared to anti-EtpA mouse polyclonal IgG, and negative IgG isotype control (-IgG) *=0.0103, ***=0.0002. **F.** mAb inhibition of blood group A-EtpA interaction **=0.0084 (Kruskal-Wallis). **G**. Anti-EtpA mAbs inhibit ETEC bacterial adhesion. Data reflect replicate experiments (n=45 fields total) and the impact of anti-EtpA mAbs on ETEC adhesion to target blood group A expressing HT-29 cells ****<0.0001, **<0.001(Kruskal-Wallis). Grey bars throughout represent geometric mean values. **H**. mAb recognition of full-length rEtpA (blue bars) and NTS of EtpA (grey bars) in end-point ELISA. Data represent geometric mean of ≥ 6 technical replicates combined from 2 experimental replicates. ****<0.0001 by ANOVA. **I.** mAb inhibition of EtpA binding to human A blood group in kinetic ELISA assay. N=8 technical replicates from 2 independent experiments (Ø=no antibody control; *<0.05, **<0.005, by Kruskal-Wallis).

### Antibodies targeting EtpA repeats interrupt bgA binding and ETEC adhesion

To identify potential protective epitopes on EtpA, we examined the capacity of anti-EtpA monoclonal antibodies (mAb) to impair EtpA binding to A blood group glycans, and interrupt pathogen-host interactions. Two mAbs isolated from rEtpA-vaccinated mice, 1G05 and 1C08, both bound to EtpA with high affinity, (Supplemental figure 1A), but recognized distinct epitopes on EtpA (Supplemental figure 1 B-C). The 1G05 mAb, which recognized the CTR domain (Supplemental figure 1D) significantly inhibited interactions with blood group A (figure 1E-F) and impaired bacterial adhesion (figure 1G). Conversely, 1C08, which recognized the NTS domain (Supplemental figure 1D), exhibited no demonstrable impact on EtpA binding to target blood group A molecules or ETEC pathogen-host interactions. This pattern was also observed in monoclonals isolated from human volunteers challenged with ETEC H10407, with the three monoclonals that recognized the NTS domain (figure 1H) failing to inhibit EtpA-bgA interactions (figure 1I). Conversely the single monoclonal (1F09) that bound the CTR significantly inhibited EtpA interactions with BgA. Collectively, these data indicate that the C-terminal repeat region of EtpA is essential to ETEC virulence, and that antibodies targeting this region can effectively inhibit interactions with the host.

### The EtpA adhesin forms an elongated β-helix

To obtain the complete structure of EtpA and gain further insight into the differential activity of the 1G05 and 1C08 mAbs, we performed cryo-EM analysis of rEtpA bound to the fragment antigen binding domains (Fab) of both mAbs, resulting in reconstructions of 3.3 and 4 Å-resolution for the 1C08 and 1G05 bound complexes, respectively (Supplemental figures 2, 3). Similar to other TpsA exoproteins, the mature EtpA molecule forms an elongated and slightly twisted 3-sided parallel β-helix ~29 nm in length (figure 2A) that can be divided into amino-terminal secretion (NTS-residues 66:640) domain, and carboxy-terminal repeat (CTR – residues 641:1534) domain. The NTS domain, also referred to as the TPS domain, is highly conserved among TpsA proteins and is required for interactions with the polypeptide transport-associated (POTRA) domains of the outer membrane β-barrel transporter (TpsB).^41,55^ The NTS domain contains the only extra-helical inserts present on EtpA (figure 2B), which fold back over the exterior of the main β-helix. Unlike the closely related pectate lyase protein, which binds carbohydrate residues in the pocket created by its inserts,^56^ those of the EtpA NTS domain do not form solvent-accessible binding pockets, and their structural and/or functional importance are unknown. The NTS domain also contains the only two α-helices in the structure, H1, and H2, the latter of which creates a ~31° kink in the otherwise straight β-helix (figure 2A-B). A similar alpha helix-induced kink was observed between the NTS and functional C-terminal domains in the crystal structure of HxuA from *Haemophilus influenzae*^46^ (<u>PDB 4RM6</u>) potentially enhancing flexibility between the two domains. The structure of the EtpA NTS domain determined here aligns closely with the recently characterized crystal structure of this region (PDB <u>8CPK</u>)^57^, is similar to the published crystal structures of the N-terminal domains of related TpsA proteins including HxuA (PDB <u>4I84</u>, ^46^ <u>PDB 4RM6</u>)^42^, the HMW1 (PDB <u>20DL</u>)^43^ adhesin from *Haemophilus influenzae,* hemolysin A from *P. mirabilis* (PDB <u>3FY3</u>)^44^, and filamentous hemagglutinin from *Bordetella pertussis* (PDB <u>1RWR</u>)^41^ as illustrated in the structure-based alignment (Figure 2C).

**Figure 2.**
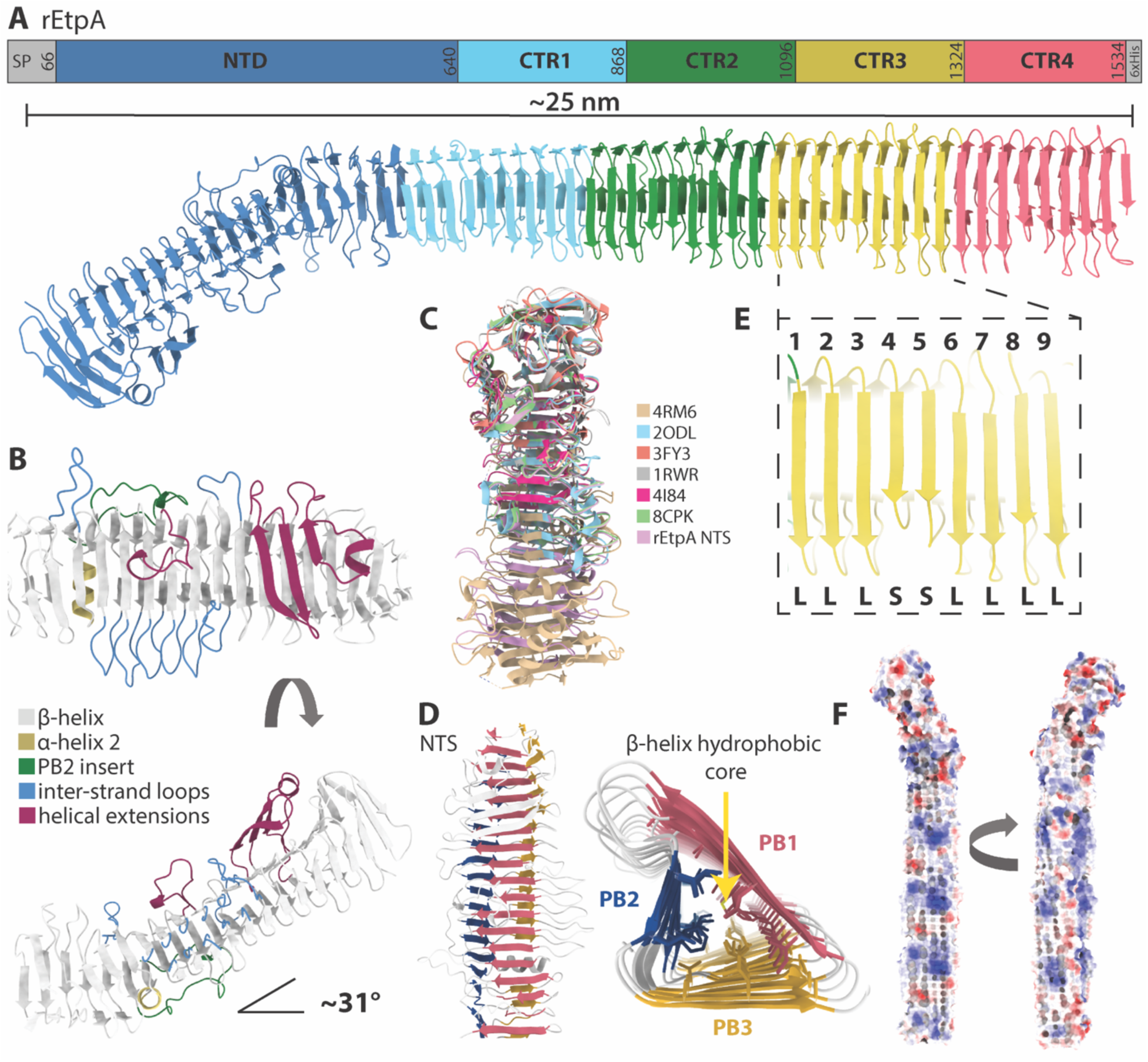
The cryo-EM structure of rEtpA. **A.** Color coded sequence diagram and atomic model of EtpA derived from cryo-EM maps. **B.** N-terminal (TPS) domain of EtpA with various features color coded. **C.** Structural alignment of previously published N-terminal TPS domains from related TpsA proteins along with their corresponding PDB identifiers. **D.** The EtpA NTD viewed from above and CTR domain viewed looking down the core of the β-helix color coded by the 3 by parallel β-sheets, PB1, PB2, and PB3. **E.** Zoomed in view of one CTR showing the 9 β-strands designated as either long (L) or short (S). **F.** Surface representation of the EtpA structure colored by Coulombic potential.

The CTR domain contains three 228 residue repeats followed by a 219-residue partial repeat forming an unbroken 3-sided β-helix (figure 2A). Following previously established convention, the three parallel β-sheets forming the sides of the helix are referred to as PB1, PB2, and PB3 (figure 2D). PB1 and PB2 are both continuous β-sheets composed of ~54 strands, while PB3 is split into two parts by H2 (figure 2A-B). Within the CTR domain, PB1 is the widest with strands that are 5-7 residues long, followed by PB3 with strands that are 4-5 residues long, then PB2 with strands that are only 2-3 residues long (figure 2D). Each CTR is composed of 9 β-strands per side (figure 2E) and separating each strand are loops which form the edges of the helix, with the loops separating PB1 and PB2 being the longest (figure 2D). The fourth and fifth strand of each repeat on PB1 are shorter than the others (figure 2E), however, the reason for this minor asymmetry is not apparent. The interior of the β-helix is composed almost entirely of closely packed hydrophobic residues which contribute to the stability of the helical structure (figure 2D), while the exterior has an abundance of polar and charged residues (figure 2F). Altogether, the lack of extra-helical extensions or apparent binding clefts on the CTR suggest that bgA must interact directly with the β-helix of this domain.

### Glycosylation of EtpA by EtpC, a promiscuous low-fidelity N-linked glycosyltransferase

Perhaps the most striking feature of the EtpA structure is its unique and extensive surface glycosylation. In addition to the *etpBAC* operon, other TPS loci from *Yersinia*, and *Burkholderia spp.* appear to encode glycosyltransferases related to HMW1C of *H. influenzae*^58–60^. The structure of the EtpC glycosyltransferase predicted by AlphaFold2^61^ shows high structural similarity (pruned Ca-RMSD = 1.1Å; all residue Ca-RMSD = 3.7Å) to the crystal structure of the closely related HMW1C glycosyltransferase from *Actinobacillus pleuropneumoniae*^62^ (Supplemental figure 4), a functional homolog to the HMW1C glycosyltransferase of *H. Influenzae* which shares ~40% sequence identity, and 56% similarity to EtpC. The HMW1 adhesin glycosylated by the HMW1C enzyme in *H. influenzae* exhibits a unique glycosylation profile consisting of asparagine-linked (N-linked) mono- and di-hexose glycans appearing predominately at canonical N-X-S/T sequences (where X is any amino acid expect proline), with a single modification of a non-canonical asparagine (Asn) residue^63^. Additional analysis identified the hexose residues as primarily glucose and sometimes galactose^59^, with the HMW1 glycosyltransferase catalyzing the formation of both the Asn-hexose and hexose-hexose linkages. HMW1C exhibited no apparent selection for modification of distinct sequons with either mono or dihexose sugars. Glycosylation by HMWC1 likely stabilizes the HMW1A adhesin and is required to tether this molecule to the surface of *H. influenzae.* Similarly, EtpC is required efficient secretion and function of the EtpA exoprotein adhesin^17,64^. Presently however, neither the glycosylation profile conferred by EtpC or its precise impact on pathogen-host interactions are understood.

To identify the location and type of glycan modifications on rEtpA we employed high-resolution site-specific mass spectrometry (MS) ^65^. We analyzed potential glycosylation at 166 out of the 196 non-tandem Asn residues (we did not detect peptides associated with 30 Asn residues), of which 96 adhere to the canonical (N-X-S/T) N-linked glycosylation sequon, and we found evidence for hexose modification at 133 sites, 94 of which meet the stricter criteria of >=25% occupancy (figure 3A). Based on the occupancy across all potential N-glycosylation sites (PNGS), mature EtpA would have on average 61 glycans per molecule, meaning that ~1 in every 24 residues (~4 %) of the EtpA exoprotein harbors an N-glycan modification. Comparatively, ≤ 2% of HMW1A^66^ and 1.7% of the SARS-CoV2 spike protein^67^ are glycosylated. Among sites with the highest occupancy (≥ 75%) only 4 out of the 29 are non-canonical Asn residues, (Supplemental figure 5), suggesting that like HMW1A, Asn residues within canonical sequons are glycosylated with higher fidelity. All 4 of these non-canonical glycosylation sites fall within the same repeating sequence/structural motif located on the last β-strand of each CTR in PB1. Though the majority of PNGS were found to be occupied with monohexose, dihexose was observed at 83 sites, but only 4 of those sites, N744, N972, N1200 and N1428, were found to have ≥ 50% dihexose, again all belonging to a common repeating structural motif located on the second short β-strand of each CTR in PB1 (figure 3A-B). Altogether these data suggest that the surface glycan coat of EtpA is both dense and variable.

**Figure 3.**
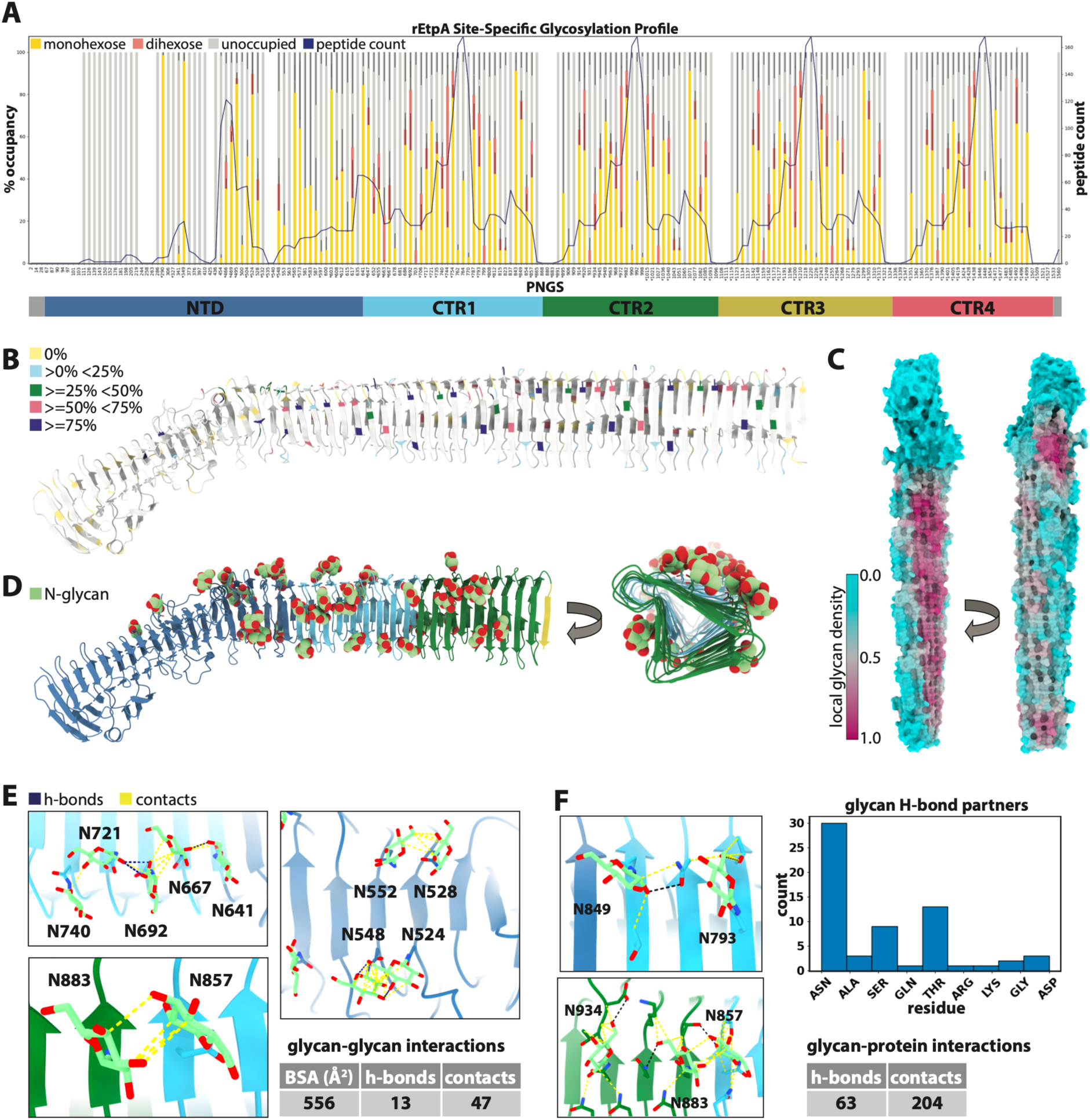
N-linked glycosylation of rEtpA. **A.** Site-specific mass-spectrometry data for rEtpA showing the % occupancy by monohexose, or dihexose at all 196 asparagine residues analyzed along with the peptide count (number of peptides detected in the experiment) and corresponding domain diagram. The * under certain residues indicates a canonical PNGS sequon site. **B.** Structure of rEtpA with potential N glycosylation sites (PNGS) colored by % occupancy. **C.** Surface representation of rEtpA structure colored by local glycan density (20Å radius). **D.** Structure of rEtpA colored by domain showing all modeled N-linked glucose residues (through CTR2) viewed from the side and looking down the core of the β-helix. **E.** Analysis of glycan-glycan interactions and (**F**) glycan-protein interactions from the atomic model of rEtpA showing hydrogen bonds (blue) and contacts (yellow) along with summary tables and a histogram of all glycan-protein residue hydrogen bonding partners.

When viewed in a structural context, we see that the confirmed N-linked glycosylation sites (NGS) on EtpA are asymmetrically distributed across the protein (figure 3B-D). PNGS within the NTS domain are glycosylated with significantly higher fidelity and specificity (for the canonical N-linked glycosylation sequon) that Asn residues within the CTR domain. For example, the NTS accounts for ~39% of the protein (and ~39% of Asn residues), however, 22 out of the 33 PNGS (~67%) without any detected glycan modifications were within the NTS, and 19 of those were within the first ~400 residues. Further, the NTS domain contains the two NGS (N290 and N349) with the highest occupancies (≥ 95%). These data suggest that the EtpA sequence as well as structural determinants may dictate glycosylation by EtpC.

### N-linked glycan clustering and intramolecular interactions on rEtpA

As illustrated by mapping the occupancy-weighted local glycan density onto the protein surface (figure 3C), the NGS are more evenly dispersed across the CTRs, but significantly more abundant on the PB1 face of the β-helix (figure 3C-D). Our cryo-EM maps confirm the location of many of these hexose modifications (figure 3D). Of the glycosylated residues up through CTR2 where the cryo-EM map permitted identification (84 >0%, 60 ≥25%, 43 ≥50%, 19 ≥75% occupancy), we were able to model hexose residues at 39. Although previous analytical studies of HMW1A reveal a mixture of glucose and galactose residues, we are unable to differentiate between glucose and galactose with MS alone, so all hexose residues were modeled as glucose for consistency (figure 3D). We did not observe clear map density for dihexose modifications at any NGS.

Given the extensive glycosylation of EtpA and the poor yields obtained in prior attempts to express the exoprotein without EtpC, we questioned whether the glycans might contribute to stabilization, folding and secretion of the protein. The stabilizing effect of N-linked glycans is at least partially mediated through favorable interactions with neighboring amino acid side chains, often via stacking with aromatic residues or hydrogen bonding with polar residues. This stabilizing interaction almost always involves the core *N*-acetylglucosamine, which in the case of EtpA would be equivalent to the N-linked hexose residue^68,69^. Although we did not find statistical enrichment of any aromatic residues around the NGS (Supplemental figure 6A-C), our structure did reveal numerous glycan-glycan, as well as glycan-amino acid interactions with other residue types (figure 3E,F). Further, the NGS on EtpA have a tendency group into local clusters. For example, of the 11 glycans modeled on PB1, 5 of them, including 1 more from the final strand of the NTS domain, are located immediately adjacent to each other on the first residue of each β-strand and can be seen to form a chain of inter-glycan interactions (figure 3E, top left). Other clusters of 2 or more glycans are observed throughout the structure on both the NTS and CTR domains (figure 3E; bottom left, right). In total, there are 13 potential glycan-glycan hydrogen bonds and 47 contacts captured in our structure, resulting in 556Å of buried glycan surface area. In addition, we identified 63 potential glycan-amino acid hydrogen bonds, with the majority involving N residues as well as T and S (Figure 3F). Although individually weak, these numerous small interactions likely contribute collectively to overall EtpA stability.

Finally, we sought to determine the glycosylation profile of native full-length EtpA from ETEC strain H10407. Despite inherent difficulties in protein purification and low peptide detection in MS, we were able to map the glycosylation state at 48 PNGS and observed close agreement with rEtpA, validating recapitulation of the glycosylation profile of the native protein by the recombinant expression system. (Supplemental figure 7).

### Molecular Interactions with the mAbs1C08 and 1G05

Our cryo-EM maps confirm that 1C08 monoclonal binds the NTS domain (figure 4A, Supplemental figure 2) while the neutralizing monoclonal 1G05 binds the CTR region (figure 4a, Supplemental figure 3). The 1G05 epitope encompasses PB1 and the inter-strand loops between PB1 and PB2 and is located at the interface of the repeat domains, allowing for the binding of up to 3 Fabs per molecule (figure 4A, Supplemental figure 4). The 1C08 epitope encompasses PB3 and the long inter-strand loops between PB3 and PB1, just before the H2 helix (figure 4A). 1G05 and 1C08 share ~88% sequence identity and have similar structures (Cα-RMSD = 0.77Å, figure 4B-C), despite being derived from different germline genes (Supplemental figure 8). Both Fabs utilize their heavy chain (HC) complementarity determining region (CDR) loops extensively, while 1C08 also makes numerous contacts with its light chain (LC) CDR loops, and their binding interfaces bury 864Å^2^ and 1275Å^2^ of surface area, respectively (figure 4D). The larger buried surface areas and more extensive interactions of 1C08 are consistent with the kinetics data showing tighter binding to rEtpA. The weaker binding of 1G05 is compensated for by avidity affects arising from the possible binding of both Fab arms of a single IgG to each EtpA molecule. Consistent with the different binding modes, analysis of somatic hypermutation from the predicted unmutated common ancestor (UCA) germline sequence using ARMADiLLO^70^ shows that 1G05 harbors 11 mutations in its HC as opposed to 7 for 1C08, while 1C08 has a more mutated LC with 6 mutations compared to 1G05 with only 3 (Supplemental figure 8).

**Figure 4.**
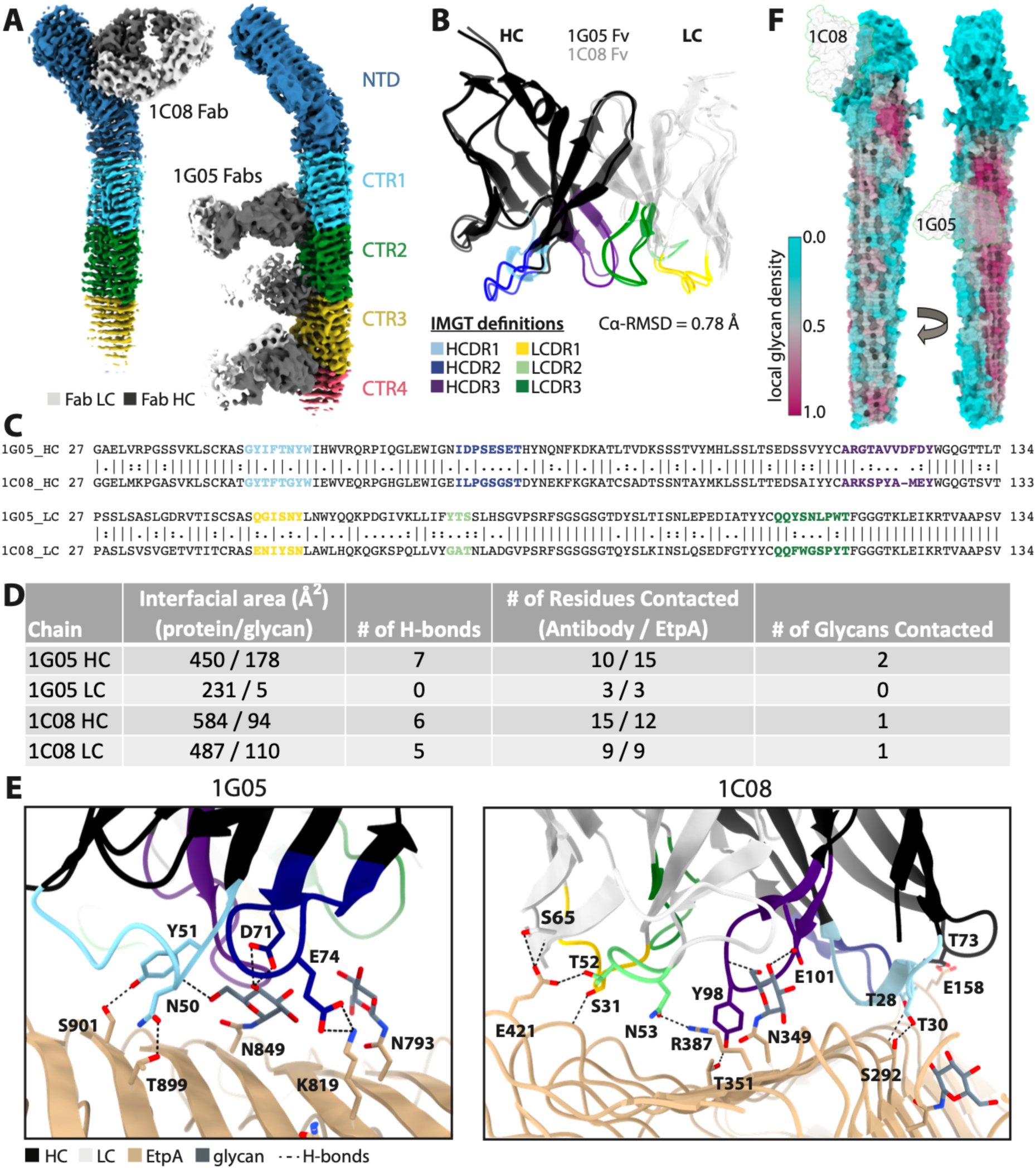
Molecular interactions between rEtpA and mAbs 1C08 and 1G05. **A.** Fabs Cryo-EM density maps of rEtpA in complex with 1C08 and 1G05 Fabs colored by domain. **B.** Structrual alignment of mAb Fc domains colored by heavy chain (HC) and light chain (LC) complementary-determining-region loops (CDRL) along with a Ca-RMSD. **C.** Pairwise sequence alignment of both Fv domains with CDRLs color coded. **D.** Table summarizing intermolecular interactions between rEtpA and mAbs calculated from PDBePISA. **E.** Intermolecular interactions between rEtpA and mAbs. **F.** Fabs 1 C08 and 1G05 projected onto glycan density maps of EtpA.

A defining feature of both epitopes is the presence of a single centrally located N-linked hexose residue, N349 and N849 for the 1C08 and 1G05 epitopes, respectively (figure 4E). Intriguingly, N349 has the second highest occupancy of any site as determined by MS (~96%) and N849 is in the top 17% of sites by occupancy. This could be taken as evidence that antibodies targeting PNGS with higher occupancy are enriched during affinity maturation, or conversely, that heterogeneity in glycosylation is being exploited as a mechanism of immune evasion. 1C08 targets an epitope with relatively low local glycan density in the NTS, while 1G05 targets an epitope with high local glycan density, however, 1G05 is oriented perpendicular to the long axis of the β-helix and utilizes a smaller HC dominant binding interface such that it positions its LC away from the heavily glycosylated surface of PB1, thus avoiding all but a single glycan residue. The local density analysis also reveals that it would be difficult for antibodies to target an entirely glycan-free epitope on the CTR domain, while there is ample glycan-free surface area for potential antibody binding on the NTD.

Lastly, to determine the extent to which these glycans contribute to the affinity of mAb 1G05 for epitopes within the CTR, we mutated EtpA asparagine residues at N849, N1077, and N1305 to alanine. Affinity of the 1G05 mAb for the mutant protein was significantly diminished while binding of 1C08 was unimpaired (Supplemental figure 9A-B). Importantly however, these mutations within the CTR did not impact interaction with target A blood group glycans (Supplemental figure 9C).

### EMPEM of sera from rEtpA vaccinated mice

Earlier studies have demonstrated that vaccination with rEtpA affords significant protection against intestinal colonization by ETEC^71–73^. To further understand the potential immunogenic landscape of rEtpA we performed negative-stain electron microscopy (NSEM)-based polyclonal epitope mapping of sera from mice vaccinated with rEtpA (figure 5A, Supplemental figure 10). We found that polyclonal antibodies target a variety of epitopes on both the CTR and NTS domains of rEtpA (figure 5B). With NSEM images of Fabs bound to rEtpA informed by the high-resolution cryo-EM structures, we were able to confidently identify antibodies bound at either the N- or C-terminal domains and estimate the total number of unique epitopes on each. We found that 5-6 and 3-4 unique epitopes are targeted on the NTS and CTR domains, respectively, suggesting that the polyclonal response to EtpA likely generates a variety of both neutralizing and decoy epitopes.

**Figure 5.**
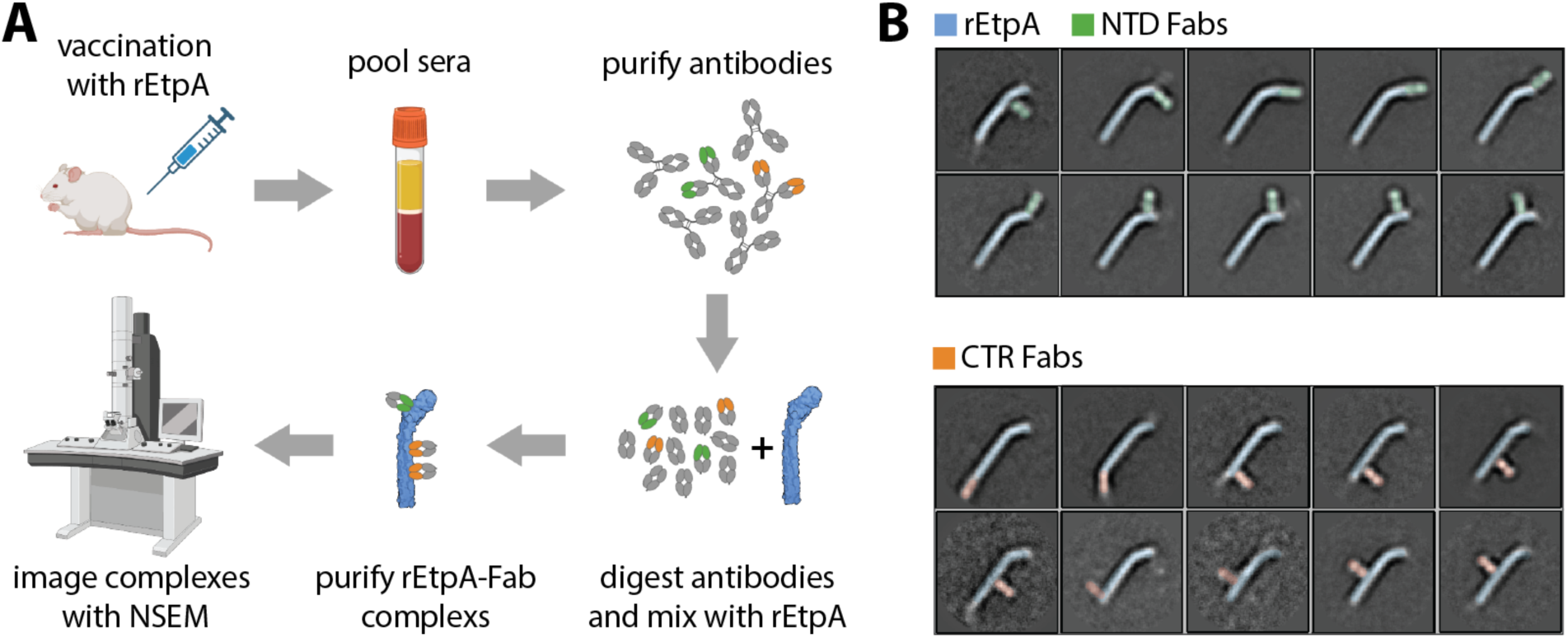
EMPEM of pooled sera isolated from rEtpA vaccinated mice. **A.** Simplified schematic of the EMPEM workflow. **B.** 2-dimensional class averages of rEtpA in complex with polyclonal Fabs showing all unique epitopes identified with EMPEM along with cartoon representations of each complex.

## Discussion

Although ETEC is an extraordinarily common cause of diarrheal morbidity in LMICs, there is presently no licensed vaccine to protect against these pathogens. To date, virtually all vaccine development efforts for ETEC have focused on a subset of heterogeneous canonical antigens known as colonization factors (CFs). To date 29 antigenically distinct CFs have been identified, potentially confounding the development of a broadly protective vaccine^74^. However, recent studies have suggested that other antigens including the EtpA adhesin are important for ETEC molecular pathogenesis, are recognized following both experimental human challenge as well as natural infections and are more highly conserved within the ETEC pathovar^75^. The studies outlined here provide a detailed structure of this complex extracellular glycoprotein adhesin molecule and offer further insight into the nature of its interactions with human host intestinal epithelia.

Clinical presentations associated with ETEC range from mildly symptomatic illness to severe diarrhea indistinguishable from cholera. Indeed, the early identification of the ETEC pathovar resulted from investigating cases of *Vibrio cholerae*-negative cholera^76^. However, in contrast to *Vibrio cholerae* infections which tend to be more severe in blood group O subjects, A blood group not O, is associated with more severe ETEC illness. Notably, toxin delivery by ETEC requires intimate cell contact ^77^. EtpA, a blood group A lectin originally identified in H10407, an ETEC isolate from a case of severe cholera-like diarrhea^78^, accelerates binding and intoxication of blood group A epithelia^79^. These studies provided a molecular basis for the enhanced disease severity among A blood group individuals observed in both young children in endemic areas^80^ as well as adult human volunteers challenged^79^ with H10407.

The present studies further define the nature of EtpA-mediated interactions with A blood group glycans. EtpA-targeting of blood group A glycans, which have N-acetylgalactosamine as their terminal sugar moiety, involves repeat modules comprising the C-terminal 2/3rds of the molecule. Interestingly, tandem repeats of carbohydrate binding modules (CBMs) ^81–83^ including those identified in other bacterial virulence molecules achieve tight binding through multivalent interactions with target host glycans^83^. The identification of epitopes critical for EtpA host interaction within the heavily glycosylated CBM repeats may have significant implications for the ETEC vaccine design. Multi-epitope fusion vaccines have been heralded as a potential strategy to achieve valency sufficient to protect against these diverse pathogens. The present studies suggest, however, that similar to HIV^84^, a thorough understanding of antigenic structure, including glycosylation profiles, combined with precise identification of protective epitopes is essential to inform rational design of immunogens^85^ that afford broad protection against these common pathogens.

The characterization of EtpC as a low-fidelity N-linked glycosyltransferase has important implications for understanding the immune response to these pathogens during infection and vaccination. Comparison of EtpA sequences from disparate geographic origins collected over time has revealed little amino acid variation despite the significant immunogenicity of this molecule^75^. The glycan-centered epitopes of mAbs characterized here may indicate that ETEC exploit glycosylation as a potential immune evasion mechanism. Many enveloped viruses hijack eukaryotic host cellular machinery to decorate their surface fusion proteins with “self” N-linked glycans facilitating immune evasion ^84, 86^. However, unlike EtpA, these viral fusion proteins can tolerate extensive mutation, allowing for PNGS to be added or removed readily. Thus, the EtpA adhesin may have evolved with its own low-fidelity glycosyltransferase under selective pressure to shield important and highly conserved functional regions of the molecule by generating a variety of N-linked glycan-profiles in immune evasion.

Negative stain EMPEM reveals that rEtpA possesses a large immunogenic surface, with multiple potential epitopes on both the NTD and CTR domain, and analysis of anti-EtpA mAbs to date indicate that only some of these antibodies, particularly those targeting the CTR domain are likely to afford protection. Given the promise of recombinant subunit-based vaccines based on these glycosylated adhesins, it will be vitally important to ensure that such vaccine antigens present native-like epitopes with the proper glycosylation.

## Acknowledgments

Anyone who contributed to the research or manuscript, but who is not a listed author, should be acknowledged (with their permission).

## Author Contributions

Z. Berndsen, J. Fleckenstein, A. Dell, S. Haslam, A. Ellebedy, J. Paulson, J. Yates, and A. Ward designed the research; Z. Berndsen, M. Akhtar, M. Thapa, T. Vickers, A. Schmitz, J. Torres, S. Baboo, P. Kumar, N. Khatoom, M. Hamrick, J. Diedrich, S. Martínez-Bartolomé, J. Turner, R. Laird, F. Poly, and C. Porter, performed experiments; Z. Berndsen, J. Fleckenstein, A. Ellebedy, and A. Ward analyzed data and wrote the manuscript. All authors revised the manuscript and approved the final version.

The views expressed in this article reflect the results of research conducted by the authors and do not necessarily reflect the official policy or position of the Department of the Navy, Department of Defense, nor the United States Government. R.L., F.P. and C.P. are federal employees of the United States government. This work was prepared as part of official duties. Title 17 U.S.C. 105 provides that ‘copyright protection under this title is not available for any work of the United States Government.’ Title 17 U.S.C. 101 defines a U.S. Government work as work prepared by a military service member or employee of the U.S. Government as part of that person’s official duties.”

## Data Availability

Cryo-EM maps and refined atomic models will be submitted to the Electron Microscopy Databank (EMDB) and Protein Data Bank (PDB) prior to publication.

## Materials and Methods

### Molecular cloning of EtpA mutants

To construct a plasmid encoding EtpA_1-1806_, a Tn7-based GPS4 transprimer-mutagenized (NEB) etpA genes^36^ with insertions in the C-terminal repeat region of *etpA* were amplified from the indicated template plasmid (supplementary table 1) with primers jf051716.1 and jf082718.1 (supplementary table 2), to permit In-Fusion (TaKaRa) cloning of the resulting *etpBA* amplicon bearing a truncated *etpA* gene into pBAD/myc-His B digested with *NcoI/HindIII*, placing the EtpA truncation in-frame with the C-terminal polyhistidine tag. Plasmid pQL211 encoding the EtpB and amino terminal secretion domain of EtpA (EtpA_1-607_) was generated from pJL017 (supplemental table 1) with primers jf042114.3/jf042114.4 (supplemental table 2).

The triple mutant EtpA-N849A-N1077A-N1305A was created by excision of the highly repetitive C-terminal region from the *etpA* gene by digestion of pJL017 with *Kfl*I and *Hind*III, then the insertion of three overlapping gene blocks f1-N849A, f2-N1077A, and f4-N1305A (IDT) by HiFi assembly (NEB). When inserted into the digested plasmid, these gene blocks created a synthetic gene with the three desired mutations. The resulting plasmid, <u>pTV005</u> was verified by long-read nanopore sequencing (<u>plasmidsaurus</u>).

### Purification of recombinant EtpA and EtpA mutants

EtpA and subclones were purified as previously described^87^. Briefly, 75 ml of Terrific Broth containing ampicillin (100 µl/ml) and chloramphenicol (15 µg/ml) with 0.2% (w/v) glucose was inoculated with frozen glycerol stock of jf1696, jf3013, or jf5090, and grown overnight at 37 degrees C, 225 rpm. 5 ml of overnight growth was used to inoculate 2-liter flasks each containing 500 ml fresh media, and grown at 37 degrees C, 225 rpm to OD ~0.6 Cultures were then induced with 0.0002 % arabinose (5 µl 20 % arabinose/flask) x 4-5 hours @ 37 degrees C, 225 rpm. Cultures were then centrifuges at 8,000 rpm x 10 min, and supernatant stored overnight at 4 degrees C. After filtration through 0.2 µm filters, the filtrate was concentrated via tangential flow (Pellicon 2 Biomax 100 kDa MWCO) to a final volume of ~100 ml and applied to a 10 ml His-Trap column, then washed with 5 column volumes of binding buffer (50 mM PO4 pH 7.5 300 mM NaCl). Protein was eluted over a gradient of 50 mM PO4, pH 7.5, 300 mM NaCl, 1 M imidazole, fractions collected and analyzed by SDS-PAGE. Fractions with protein of interest were pooled and dialyzed vs 10 mM MES pH 6, 100 mM NaCl, then concentrated to final concentration of ~1 mg/ml.

For purification of native EtpA (nEtpA) from H10407, the flagellin null mutant (<u>jf3099</u>) was grown in 4 L LB media and the culture supernatant obtained and concentrated as before. The concentrate was then exchanged into 50 mM MES pH 6, 300 mM NaCl, 1 mM EDTA and proteins precipitated by addition of ammonium sulfate to 80% saturation. The resulting precipitate was recovered by centrifugation at 9,820 xg for 10 min, and the pellet redissolved in 5 ml PBS. This solution was then concentrated to 1 ml using a 50 K cutoff spin concentrator (Millipore). The sample was applied to a HiLoad 16/600 superdex 200 gel filtration column equilibrated in PBS, and proteins separated at a flow rate of 1 ml min^-1^. Fractions containing nEtpA were identified by SDS-PAGE, pooled, and concentrated to produce the final preparation.

### EtpA biotinylation

Primary amines of recombinant EtpA were biotinylated with EZ-Link Sulfo-NHS-LC-LC-Biotin (ThermoFisher <u>21388</u>) for 30 minutes at room temperature. The reaction was quenched with Tris 100 mM, pH 8.0, then reactants dialyzed to remove excess biotin.

### Mouse immunization and plasmablast sorting

All procedures involving animals were performed in accordance with guidelines of Institutional Animal Care and Use Committee of Washington University in Saint Louis (protocol number 21-0053). Two female C57BL/6J mice (Jackson Laboratories) were immunized intramuscularly and boosted nine weeks later with 15 µg EtpA emulsified in PBS and AddaVax (InvivoGen). Draining iliac and inguinal lymph nodes were collected four days after boosting and single cell suspensions were prepared for plasmablast sorting. Cells were stained for 30 min on ice with CD138-BV421 (281-2, 1:200), IgD-FITC (11-26c.2a, 1:100), CD19-PE (1D3, 1:200), CD38-PE-Cy7 (90, 1:200), Fas-APC (SA367H8, 1:400), CD3-APC-Cy7 (17A2, 1:100), and Zombie Aqua (Biolegend) diluted in PBS supplemented with 2% FBS and 2mM EDTA. Cells were washed twice and single PBs (CD138^+^ CD38^lo^ CD19^+/lo^ CD3^-^ live singlet lymphocytes) were sorted using a FACSAria II into 96-well plates containing 2 µL Lysis Buffer (Clontech) supplemented with 1 U/µL rNase inhibitor (NEB) and immediately frozen on dry ice.

### Plasmablast isolation from volunteers experimentally challenged with ETEC

Peripheral blood mononuclear cells obtained from human volunteers challenged with ETEC strain H10407^88^ were obtained from archived specimens maintained at the National Medical Research Command, Silver Spring, Maryland. The study protocol was approved by the Naval Medical Research Command Institutional Review Board in compliance with all applicable federal regulations governing the protection of human subjects. Use of de-identified specimens from volunteers was approved by the Washington University in Saint Louis Institutional Review Board (Protocol number 201110126). Cells were incubated for 10 min on ice with FcX (Biolegend), then stained for 30 min on ice with biotinylated rEtpA in PBS supplemented with 2% FBS and 2mM EDTA. Cells were washed twice and stained for 30 min on ice with CD4-Spark UV 387 (SK3, 1:200), CD8-Spark UV 387 (SK1, 1:100), CD14-Spark UV 387 (63D3, 1:200), CD20-Pacific Blue (2H7, 1:400), IgD-BV785 (IA6-2, 1:200), CD19-FITC (HIB19, 1:100), CD71-PE (CY1G4, 1:400), CXCR5-PE-Dazzle594 (J252D4, 1:50), CD38-PE-Fire810 (S17015F, 1:400), streptavidin-APC, and Zombie NIR (all Biolegend) diluted in PBS supplemented with 2% FBS and 2mM EDTA. Cells were washed twice and single rEtpA-binding PBs (rEtpA^+^ CXCR5^low^ CD71^+^ CD20^low^ CD38^+^ IgD^low^ CD19^+/int^ CD4^−^ CD8^−^ CD14^−^ live singlet lymphocytes) were sorted using a Bigfoot into 96-well plates containing 2 µL Lysis Buffer (Clontech) supplemented with 1 U/µL rNase inhibitor (NEB) and immediately frozen on dry ice.

### Monoclonal antibody isolation and purification

Antibodies were cloned as previously described^89,90^. Briefly, VH and Vκ genes were amplified by reverse transcriptase-polymerase chain reaction (RT-PCR) and nested PCR from single-sorted plasmablasts using cocktails of primers specific for IgG, IgM/A, Igκ, and Igλ using first round and nested primer sets^89–92^ and then sequenced. Clonally related cells were identified by the same length and composition of IGHV, IGHJ and heavy-chain CDR3 and shared somatic hypermutation at the nucleotide level. To generate recombinant antibodies, heavy chain V-D-J and light chain V-J fragments were PCR-amplified from 1^st^ round PCR products with mouse variable gene forward primers and joining gene reverse primers having 5’ extensions for cloning by Gibson assembly as previously described^90,93^ and were cloned into the pABVec6W human IgG1 antibody expression vector^94^ in frame with either human IgG or IgK constant domain. Plasmids were co-transfected at a 1:2 heavy to light chain ratio into Expi293F cells using the Expifectamine 293 Expression Kit (Thermo Fisher), and antibodies were purified with protein A agarose (Invitrogen). From 94 sorted cells, 49 rearranged IGHV sequences were recovered, of which 31 were clonally distinct. 7 clonally distinct mAbs were generated, of which 5 bound EtpA.

### EtpA human A blood group interaction studies

#### Erythrocyte pull down

Erythrocyte pull-down studies were performed with 6His-tagged EtpA linked to cobalt-coated magnetic beads (Dynabeads, Invitrogen 10103D) and blood group A1 red blood cells (RBC) obtained from Immucor (0002345). Briefly, beads were re-suspended by vortexing, washed with 1 ml of PBS, then re-suspended in 100 µl of PBS and 100 µl of EtpA-6His at a final concentration of 1 mg/ml, and incubated at room temperature on a rotary mixer (~10 rpm) x 20 minutes. Beads were separated on a magnet, the supernatant removed, and then washed in 1 ml of PBS. After re-suspension in 100 µl of fresh PBS, EtpA-linked beads were maintained on ice during RBC preparation. A1 RBCs were re-suspended by inverting the tube of cells, and 1 ml of the suspension was transferred to a 1.5 ml microfuge tube on ice. RBCs were spun at 1000 rpm at 4 degrees C x 1 minute. Supernatant was then removed and RBCs were washed 4 times in low ionic strength solution (LISS)–-1.75 g/L NaCl, 18 g/L glycine, 0.01 %(w/v) sodium azide, 11.3 ml of 150 mM KH2PO4 stock, 8.7 ml of 150 mM Na2HPO4 stock, final pH 7.0

#### EtpA-blood group A ELISA

Blood group A conjugated to BSA (Dextra NGP6305) was dissolved in PBS containing 0.02 %(w/v) azide to a final concentration of 0.5 mg/ml, and stored at 4 degrees C prior to use. A working solution of bgA-BSA conjugate was prepared in carbonate buffer, pH 9.6 at a final concentration of 1 µg/ml. 100 µl of the solution was then used to coat each well of ELISA strips (Corning 2580) overnight at 4 degrees C. Wells were then washed 3 x with 200 µl of PBS containing 0.02 % Tween-20 (PBS-T), then blocked with 100 µl of 1 % BSA in PBS-T at 37 degrees C for 1 hour. Coated wells were then probed with 100 µl of EtpA-biotin (10 µg/ml in PBS-T-1 % BSA)/ well for 2 hours at room temperature. Plates were washed 5 x with of PBS-T, and incubated with avidin-HRP conjugate (BioRad 1706528, diluted 1:10,000 in 1 % BSA in PBS-T) for 1 hour at room temperature, then washed again 4x with PBS-T. Plates were developed with freshly prepared room-temperature HRP substrate (TMB-(3,5,’’,’’-tetramethylbenzidine)-2 component reagent (seracare 5120-0053), and read kinetically at 650 nm (blue).

### ETEC adhesion to blood group A intestinal epithelia

HT-29 cells (ATCC HTB-38) which express blood group A glycans were propagated as previously described in McCoy’s-5A medium (Gibco, Life Technologies) supplemented with 10 % bovine serum albumin. Cells were grown to confluence in 96-well plates and incubated at 37degrees C, 5 % CO2 for one week prior to use. Adhesion assays were performed as previously described23 using mid-log phase bacterial cultures. After 30 minutes monolayers were washed 3 x with pre-warmed media, then treated with 0.1 % Triton-X-100 in PBS for five minutes. Dilutions of the resulting lysates were plated onto Luria agar and bacterial adherence expressed as the percentage of the original inoculum recovered.

### Confocal laser scanning microscopy

HT-29 cells seeded onto poly-L-lysine treated glass coverslips were incubated in 24 well plates at 5% CO2, 37ᵒC to confluence. Bacteria were added at a multiplicity of infection of ~1:100 and incubated for 1 hour prior to fixation. CellMask deep red plasma membrane stain (Thermo Fisher Scientific, C10046) (1:2,000) and DAPI (1:6000) were used to stain cells and A blood group was detected with mouse monoclonal antibody Z2A (Santa Cruz sc-69951) against human A blood group antigen, followed by AlexaFluor 647-conjugated goat anti-mouse IgM heavy chain (Molecular Probes, A21238). Confocal images were acquired using a Nikon Eclipse Ti2 inverted microscope. ETEC H10407 (serotype O78) were imaged using polyclonal antisera (Rabbit) supplied by the Penn State E. coli Reference Center, followed by cross-absorbed goat anti-rabbit IgG (H\&\L) conjugated to either AlexaFluor 488 or 594 fluorophores (Invitrogen).

### Electron Microscopy

#### Cyro-EM sample preparation

450 μg of purified rEtpA was combined with 12x molar excess of 1G05 or 1C08 Fabs and incubated overnight at 4ᵒC. Complexes were then purified via SEC with a HiLoad 16/600 Superdex 200 pg column (GE Healthcare) on an AKTA Pure 25M system (Cytiva) using MES pH 6 as the running buffer. SEC peaks corresponding to rEtpA:Fab complexes were pooled and concentrated with Amicon 10 kDa concentrators to a final concentration of ~1 mg/ml. 3 μL of each complex was briefly incubated with lauryl maltose neopentyl glycol (LMNG; Anatrace) to a final concentration of 0.005 mM and then deposited on glow discharged Quantifoil Cu 1.2/1.3-300 mesh grids and plunge frozen using a Thermo Fisher Vitrobot Mark IV at 4ᵒC, 100% humidity, blot force 1, 10 second wait time, and a blot time of 4-7 seconds.

#### Cryo-EM–-data collection

EM micrographs of rEtpA in complex with 1G05 or 1C08 Fabs were collected on a Thermo Fisher Glacios cryo-electron microscope operated at 200keV and 96k x magnification (pixel size = 0.725-angstrom), equipped with a Thermo Fisher Falcon 4 direct electron detector, and operated with Thermo Fisher <u>EPU 2</u> software. For the rEtpA + Fab 1G05 complex, ~9,000 micrographs were collected each with a total dose of ~47 e-/angstrom-squared, fractionated over 40 frames, at a target defocus range of −1.5 to −0.5 micrometer–-For a complete description of imaging conditions and data statistics see Supplemental table 3.

#### Cryo-EM–-data processing

Both Cryo-EM datasets were processed with CryoSparc v2^95^ using the following standard workflow. Movie micrographs were aligned, and dose weighted with patch motion correction and the contrast transfer function (CTF) was fit to each micrograph using the patch-based CTF algorithm followed by manual curation to remove micrographs with poor CTF fit parameters and ice thickness. Blob picking was performed on a subset of curated micrographs followed by particle extraction, 2D-classification, and subset selection. Particles associated with good 2D classes were used to train a Topaz neural network^96^ which was then used to pick particles from the entire curated dataset. These particles were then extracted and multiple rounds of 2D classification followed by subset selection were performed. Next, multiple rounds of reference free *Ab Initio* 3D classification followed by subset selection were performed to identify particle stacks that refine well and to separate particles with different stoichiometric ratios of bound Fabs, ranging from 0-3 Fabs in the case of 1G05 and 0-1 Fabs for 1C08. These subsets were then refined separately using non-uniform 3D refinement^97^ with per-particle and global CTF refinement enabled^98^. A final round of focused 3D refinement was performed with masks around Fab epitopes.

#### Cryo-EM–-Model building and refinement

AlphaFold2^99^ implemented through <u>ColabFold</u>^100^ was used to generate the starting EtpA model for refinement. Due to the large size of EtpA the sequence was broken down into ~700 residue fragments with ~100 residue overlaps (Supplemental figure 11) and each fragment was aligned in <u>UCSF Chimera</u> ^101^ using the 100 residue overlaps followed by removal of repeated residues and model combination into a single PDB file. Fab models were generated with SABPred^102^ and added to the complete rEtpA AlphaFold2 model in <u>UCSF Chimera</u> and combined into a single PDB file. This combined file was then refined using ROSETTA^103^ asking for ~300 models. Each model was scored with MolProbidity^104^ and EMRinger^105^ and the model with the top combined score was selected. Next, N-linked glucose residues were added manually using COOT^106^ and a ligand restraint file was generated in Phenix^107^ using eLBOW^108^. Glucose residues were covalently linked to 39 PNGS up through the end of CTR2, after which map resolution diminished. The cryo-EM maps also revealed probable hexose density at 4 sites that were not observed to be glycosylated by MS, namely N540, N883, N552, and N685, the first two of which occur within peptides not detected by MS, while the last two are incorrectly assigned tandem Ns. This glycosylated model was then refined with Phenix real-space refinement and model quality was assessed with the software mentioned above. N-linked glycans were validated with Privateer^109^. If adjustments were necessary, they were performed manually in COOT and followed up with additional rounds of real-space refinement in Phenix.

### Structural Analysis

Molecular graphics for images and molecular contact calculations were performed with <u>UCSF</u> <u>ChimeraX</u>, developed by the Resource for Biocomputing, Visualization, and Informatics at the University of California, San Francisco, with support from National Institutes of Health R01-GM129325 and the Office of Cyber Infrastructure and Computational Biology, National Institute of Allergy and Infectious Diseases^110^. Epitope-Paratope interactions were analyzed with PDBePISA^111^.

### Negative Stain EM polyclonal epitope mapping

Sera obtained from CD-1 mice vaccinated intranasally (IN) with recombinant full-length EtpA - myc-His (rEtpA) were pooled and used to prepare polyclonal IgG as described previously^112,113^. Mice were vaccinated IN with 20 µg of rEtpA adjuvanted with 1 µg of dmLT^114^ on days 0, 14, 21 as previously described^71–73^, followed by terminal bleed on day 35. 5 ml of pooled mouse sera was then used to purify IgG using Protein A Sepharose resin (GE Healthcare) and digested for using papain-agarose resin (Thermo Fisher Scientific). Fc and undigested IgG were removed through with Protein A Sepharose resin using 0.2 ml packed resin per 1 mg of IgG. Fab samples were concentrated, and buffer exchanged to TBS using Amicon ultrafiltration units with a 10 kDa cutoff (EMD Millipore Sigma) and mixed in excess with rEtpA and allowed to incubate overnight at 4ᵒC. rEtpA:polyclonal Fab complexes were then purified via SEC, concentrated to ~0.01mg/ml and prepared for imaging as described previously using uranyl formate stain^112^. EM micrographs were collected on an FEI Spirit microscope operating at 120keV and controlled with Leginon software^115^. Single-particle negative stain image processing was performed with Relion/4.0 software^116^ as described previously^112^. For figures, features in 2D-class averages corresponding to Fabs were identified by eye and false-coloring was applied for clarity.

### Site-specific glycosylation analysis of rEtpA

#### Sample Preparation

Recombinant EtpA glycoprotein was exchanged to water using Microcon Ultracel PL-10 centrifugal filter. Glycoprotein was reduced with 5 mM tris(2-carboxyethyl) phosphine hydrochloride (TCEP-HCl) and alkylated with 10 mM 2-Chloroacetamide in 100 mM ammonium acetate for 20 min at room temperature (RT, 24ᵒC). Glycoprotein was digested with 1:25 Proteinase K (PK) for 30 min at 37ᵒC or 1:20 trypsin for 16 h at 37ᵒC. Proteases (PK/trypsin) were denatured by incubating at 90ᵒC for 15 min, further diluted in buffer A, subsequently analyzed by LC-MS/MS.

Proteinase K and trypsin treatment: recombinant EtpA glycoprotein was exchanged to water using Microcon Ultracel PL-10 centrifugal filter. Glycoprotein was reduced with 5 mM tris(2-carboxyethyl) phosphine hydrochloride (TCEP-HCl) and alkylated with 10 mM 2-Chloroacetamide in 100 mM ammonium acetate for 20 min at room temperature (RT, 24ᵒC). Glycoprotein was digested with 1:25 Proteinase K (PK) for 30 min at 37ᵒC or 1:20 trypsin for 16 h at 37ᵒC. Proteases (PK/trypsin) were denatured by incubating at 90ᵒC for 15 min, further diluted in buffer A, subsequently analyzed by LC-MS/MS

#### LC-MS/MS

Samples were analyzed on an Orbitrap Eclipse Tribrid mass spectrometer. Samples were injected directly onto a 25 cm, 100 μm ID column packed with BEH 1.7 μm C18 resin. Samples were separated at a flow rate of 300 nL/min on an EASY-nLC 1200 UHPLC. Buffers A and B were 0.1% formic acid in 5% and 80% acetonitrile, respectively. The following gradient was used: 1– 25% B over 100 min, an increase to 40% B over 20 min, an increase to 90% B over another 10 min and held for 10 min at 90% B for a total run time of 140 min. Column was re-equilibrated with buffer A prior to the injection of sample. Peptides were eluted from the tip of the column and nanosprayed directly into the mass spectrometer by application of 2.8 kV at the back of the column. The mass spectrometer was operated in a data dependent mode. Full MS1 scans were collected in the Orbitrap at 120,000 resolution. The cycle time was set to 3 s, and within this 3 s the most abundant ions per scan were selected for HCD MS/MS at 35 NCE. Dynamic exclusion was enabled with exclusion duration of 60 s and singly charged ions were excluded.

### Data processing

Protein and peptide identification were done with Integrated Proteomics Pipeline (IP2). Tandem mass spectra were extracted from raw files using RawConverter^117^ and searched with ProLuCID^118^ against a database comprising UniProt reviewed (Swiss-Prot) proteome for *Escherichia coli* K12 (UP000000625) with amino acid sequence for EtpA (NCBI: WP001080112.1) containing C-term MYC and 6xHis-tag, and a list of general protein contaminants. The search space included no cleavage-specificity for PK, and half-tryptic specificity with unlimited missed cleavages for trypsin. Carbamidomethylation (+57.02146 C) was considered a static modification. Monohexose (+162.052824 N) and Dihexose (+324.105647 N) were considered differential modifications and a maximum of 3 differential modifications were allowed per peptide. Data was searched with 50 ppm precursor ion tolerance and 500 ppm fragment ion tolerance. Identified proteins were filtered using DTASelect2^119^ and utilizing a target-decoy database search strategy to limit the false discovery rate to 1%, at the spectrum level^120^. A minimum of 1 peptide per protein, and no tryptic end per peptide for PK and 1 tryptic end per peptide for trypsin were required and precursor delta mass cut-off was fixed at 10 ppm. Statistical models for peptide mass modification (modstat) were applied (for trypsin, trypstat statistics used). Census2 (Park et al., 2008) label-free analysis was performed based on the precursor peak area, with a 10 ppm precursor mass tolerance and 0.1 min retention time tolerance. Data analysis using GlycoMSQuant^121^ was implemented to automate the analysis. GlycoMSQuant summed precursor peak areas across replicates, discarded peptides without NGS, discarded misidentified peptides when N-glycan remnant-mass modifications were localized to non-NGS asparagines and corrected/fixed N-glycan mislocalization where appropriate. Census output files were modified to accommodate differential mass modification notations in GlycoMSQuant. The GlycoMSQuant algorithm was modified to accommodate quantification at all asparagines without implementing sequon-specific definition of NGS.

### Sequenced-based analysis

Statistical analysis of PNGS flanking residue frequencies were performed with custom python scripts available at (https://github.com/ZTBioPhysics/EtpA-Glycosylation-Analysis.git). Briefly, for every residue in the input list, the identity of the amino acids 2 residues immediately upstream and downstream were determined and the overall frequencies for each amino acid type were calculated. To calculate p-values a permutation test was used. 1000 replicates of randomly generated residues drawn from the EtpA sequence without replacement of the same length as the original list were generated and analyzed in the same manner. The number of those replicates with higher or lower frequencies than the input list for each amino acid type were used to calculate empirical p-values. P < 0.001 was labeled with‘’**’’, p < 0.01 with’’*’’, p < 0.05 with‘’’’, and p > 0.05 with‘’n’’.

### Residue proximity analysis

Statistical analysis of neighboring residues was performed with custom python scripts available at (https://github.com/ZTBioPhysics/EtpA-Glycosylation-Analysis.git). Briefly, for every residue in the input list, the identity of all the amino acids within a given distance (here 6Å) were determined and the overall frequencies for each amino acid type were calculated. Emperical p-values were calculated with the same type of permutation test as described above.

### Local glycan density analysis

The local glycan density calculations were performed with custom python scripts available at (https://github.com/ZTBioPhysics/EtpA-Glycosylation-Analysis.git). Briefly, the EtpA atomic model was divided into 3 separate structures corresponding to each parallel β-sheet, PB1, PB2, and PB3, so that local proximity measurements would not include residues on opposite sides of the β-helix. For each residue in each of the 3 split structures, the number of PNGS withing a given distance were calculated for every residue then normalized between 0 and 1 and the B-factor column in each PDB file was replaced with this value. The three structures were then combined and the UCSFChimeraX function ‘render-by-attribute’ was used to color the surface representation of rEtpA with the local glycan density value. Considering the size of an Fab variable domain is ~40Å width-wise, a 20Å probe radius was used to simulate the local density that would be encountered by a Fab binding to the surface.

### AlphaFold2 modeling of EtpC

The sequence for EtpC (GenBank <u>AAX13510.1</u>) was fed to AlphaFold2 implemented via <u>ColabFOLD</u>^122^ with standard parameters. The top ranking model was compared to the crystal structure of HMW1C (<u>PDBID:3Q3E</u>) in <u>UCSF Chimera</u>.

### Biolayer interferometry

The binding affinities of 1C08 and 1G05 Fabs to rEtpA were measured using an Octet Red 96 instrument (Forte Bio) and Ni-NTA biosensors. The entire assay was performed in 1X kinetics buffer (1X PBS pH 7.4 + 0.02% Tween-20 + 0.1% bovine serum albumin). There was a total of five steps in the assay: baseline 1 (60 sec), loading (180 sec or 1 nm threshold, whichever came first), baseline 2 (60 sec), association (180 sec), and dissociation (600 sec). The rEtpA antigen was loaded on the Ni-NTA biosensors at 25 ug/ml and the fabs were tested at 7 different concentrations, starting at 500 nM to 7.81 nM (2-fold dilutions). Using the Octet Analysis software (ForteBio), the data were reference subtracted and curves were fit with a 1:1 binding model. K_D_, and On and off-rates were determined with a global fit.

## Supplemental information

### Statistical analysis of EtpA N-linked glycosylation

Considering the extreme heterogeneity in occupancy and the relaxed sequence preferences of the EtpC glycosyltrasferase as revealed by MS, we sought to determine if there were any statistically significant trends in the frequency of amino acid types flanking PNGS that could potentially explain this data. To do this, we calculated the frequencies of each amino acid type at the two sequence positions upstream and downstream of each glycosylated PNGS for both sequon and non-sequon sites binned into either the top or bottom half of occupancy percentage as well as those sites that did not harbor any glycan modifications (0% occupancy) (Supplemental figure 5). Looking at sequon PNGS only (Supplemental figure 5A), we see potential trends emerge. First, there is a ~3:1 bias for T over S at the N+2 position (relative to the glycosylated Asn) in all occupancy bins as well as a significant enrichment T in the N-2 position, the latter of which could be explained by the common occurrence of multiple PNGS in a row. Other trends include an enrichment of isoleucine (I) at the N+2 position among glycosylated PNGS as well as S at the N-1 position and alanine (A) at the N-2 and N+1 position among the highest occupancy sites. 7 out of the 8 sequon PNGS with 0% occupancy occur at the same sequence/structural motif, namely, at the second to last residue on the β-strand of PB3 (Supplemental figure 5B). Looking at non-sequon PNGS, one potential trend is the enrichment of the small non-polar residues glycine (G) and A at several of the flanking sites (Supplemental figure 5C). Other residues that are enriched around non-sequon sites harboring glycans are I, S, and T. Notably, there are only 13 non-sequon PNGS that have >=50% occupancy representing 3 unique sequence motifs that almost all occur at the same structural motif, namely, at the first residue of a β-strand immediately after a loop separating faces of the β-helix (Supplemental figure 5D).

### Supplemental figures

**Supplemental figure 1.**
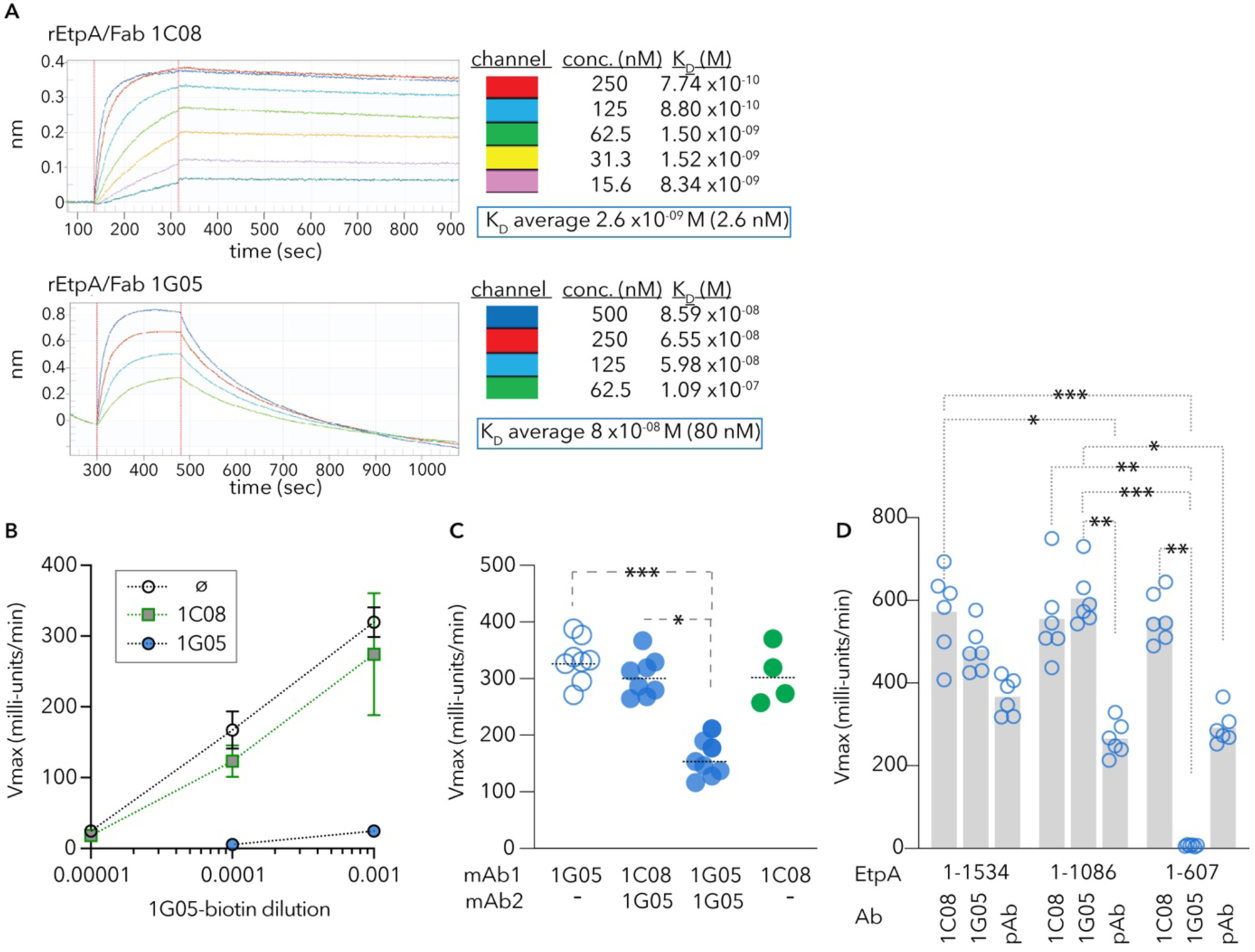
Anti-EtpA mAbs exhibit distinct affinities to unique regions of the antigen. **A**. Biolayer inferometry (Octet) studies 1C08 and 1G05 Fabs binding to rEtpA. **B.** mAb 1C08 does not compete for binding with 1G05. Shown are kinetic ELISA data indicating binding of bioinylated 1G05 mAb in the presence of unlabeled 1G05 (blue), 1C08 (green) or alone (open circle). **C.** 1G05 and 1C08 recognize EtpA but compete for different binding sites. **D.** 1G05 recognizes the repeat region of EtpA while 1C08 binds the N-terminal secretion domain. Data include n=6 technical replicates and are representative of three independent experiments. pAb=polyclonal anti-EtpA antibody. Comparisons by Kruskal-Wallis ***≤0.001, **≤0.01, *<0.05.

**Supplemental figure 2.**
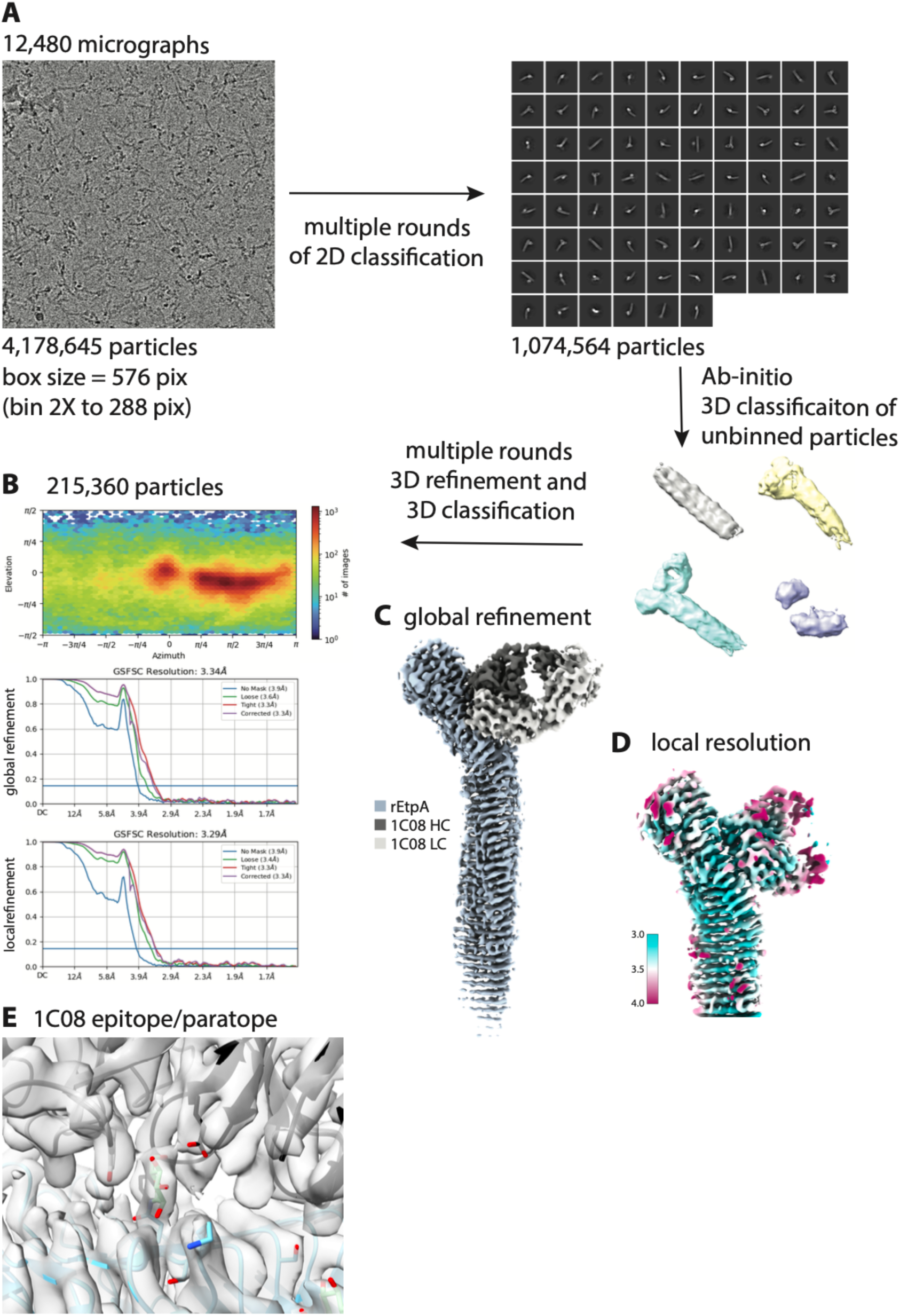
Cryo-EM data processing workflow for rEtpA-1C08 complex. Simplified cryo-EM data processing workflow including **A** representative example of an aligned and dose-weighted micrograph (lowpass filtered to 5Å), 2D and 3D class averages, and particle counts at each step. **B.** Angular distribution and Fourier shell correlation plots for the final 3D reconstruction along with the final particle count. **C.** Sharpened map colored by domain. **D.** Map colored by local resolution estimate. **E.** View of the map-model fit in the epitope/paratope region.

**Supplemental figure 3.**
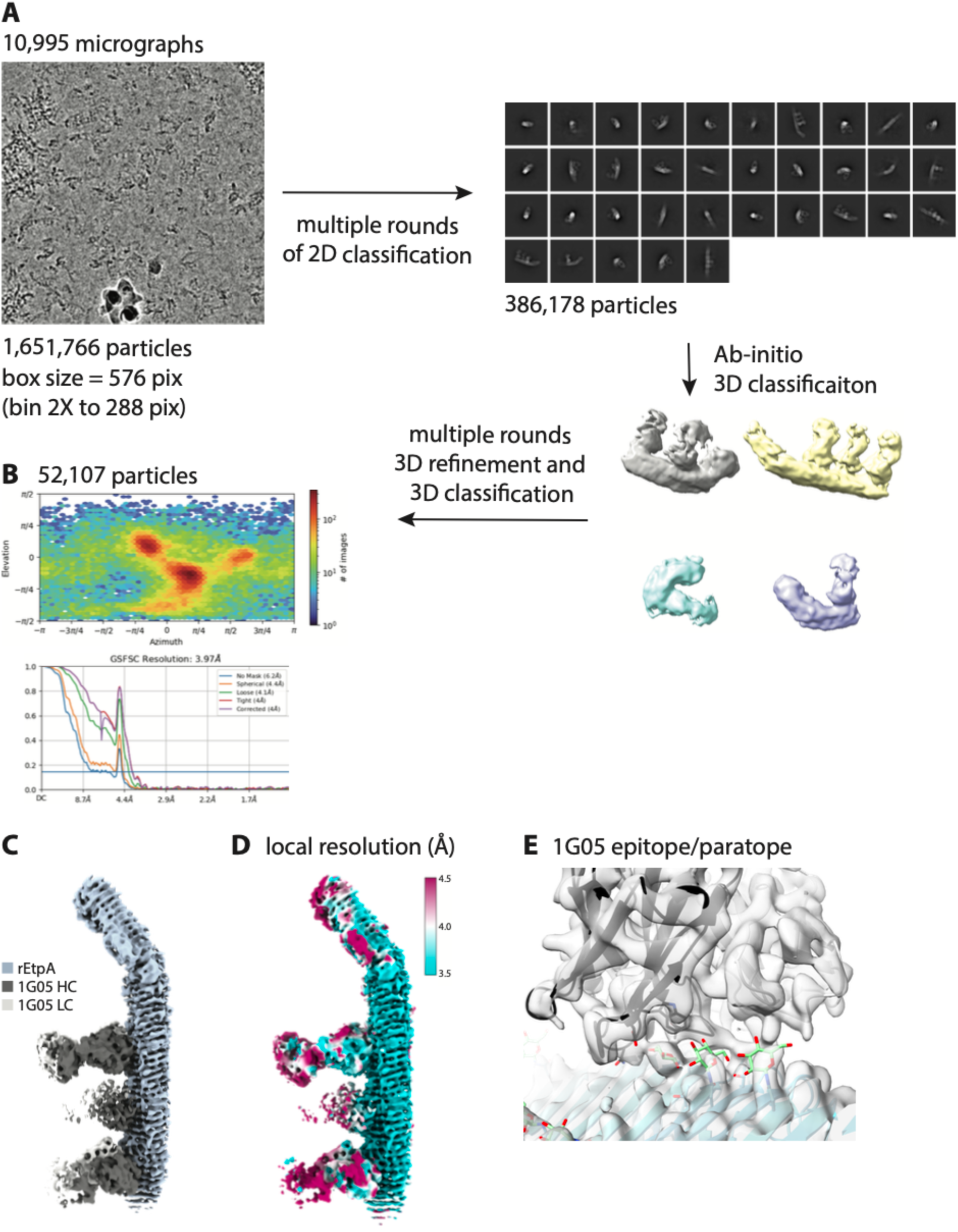
Cryo-EM data analysis for rEtpA-1G05 complex. Simplified cryo-EM data processing workflow including representative (**A**) aligned and dose-weighted micrograph (lowpass filtered to 5Å), 2D and 3D class averages, and particle counts at each step. **B.** Angular distribution and Fourier shell correlation plots for the final 3D reconstruction along with the final particle count. **C.** Sharpened map colored by domain. **D.** Map colored by local resolution estimate. **E.** View of the map-model fit in the epitope/paratope region.

**Supplemental figure 4.**
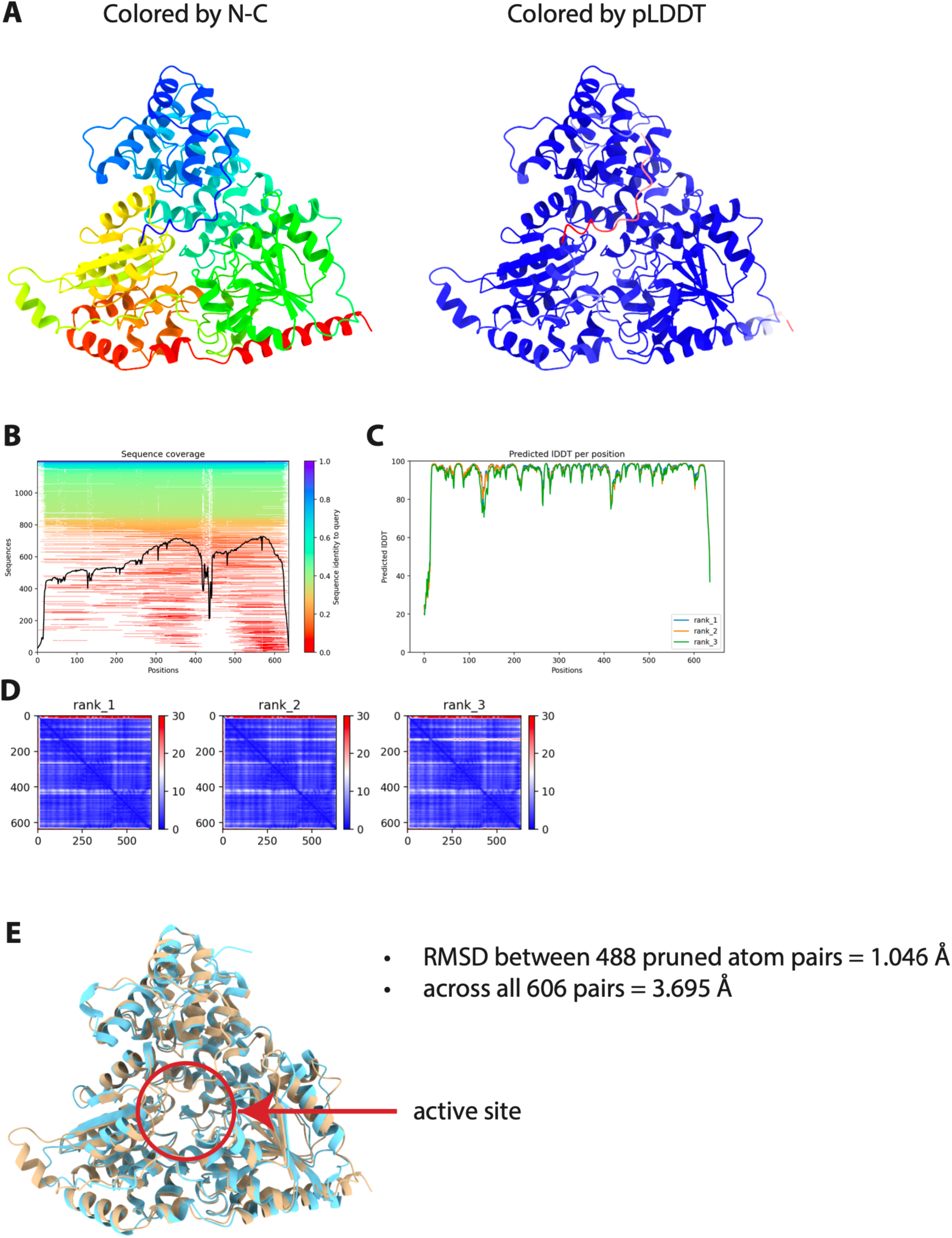
AlphaFold2 prediction of EtpC structure and structure-based alignment with the HMW1C crystal structure. **A.** Predicted structure of EtpC colored with a rainbow color mapping from the N-terminus to C-terminus and **(B)** by prediction LDDT confidence store. **C.** Multiple sequence alignment coverage. **D.** Predicted LDDT score by residue position. **E.** Predicted alignment error matricies. **F.** Structure-based alignment of predicted EtpC structure with the crystal structure of HMW1C (<u>PDBID:3Q3E</u>).

**Supplemental figure 5.**
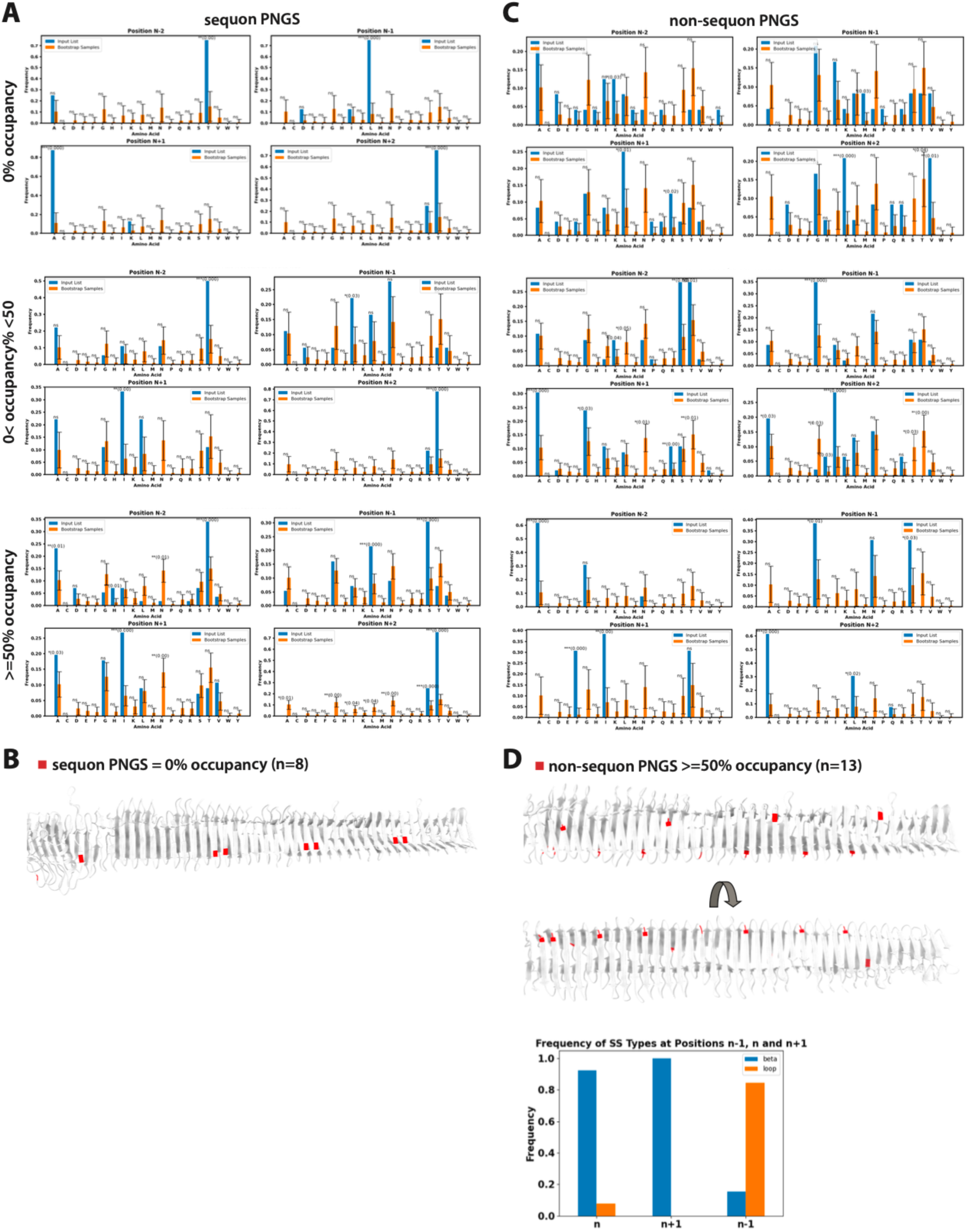
Statistical analysis of residues flanking PNGS. Statistical analysis of amino acid type frequencies at the 4 sites immediately upstream and downstream of each canonical sequon PNGS broken down by occupancy percentage as determined by mass-spectrometry. Blue bars are the average frequencies for the input residue list and orange bars are average frequencies for 1000 permutation samples with error bars and statistical significance measurements broken down by minor significance (*), median significance (**), and high significance (***) along with corresponding p-values. **B.** rEtpA structure with the 8 sequon PNGS sites with 0% occupancy colored red. **C.** Same as in A but for all non-canonical sequon PNGS. **D.** rEtpA structure showing all non-sequon PNGS with ≥ 50% occupancy colored red along with a bar plot showing the frequency of secondary structure types at the PNGS and the two residues immediately upstream and downstream of the site.

**Supplemental figure 6.**
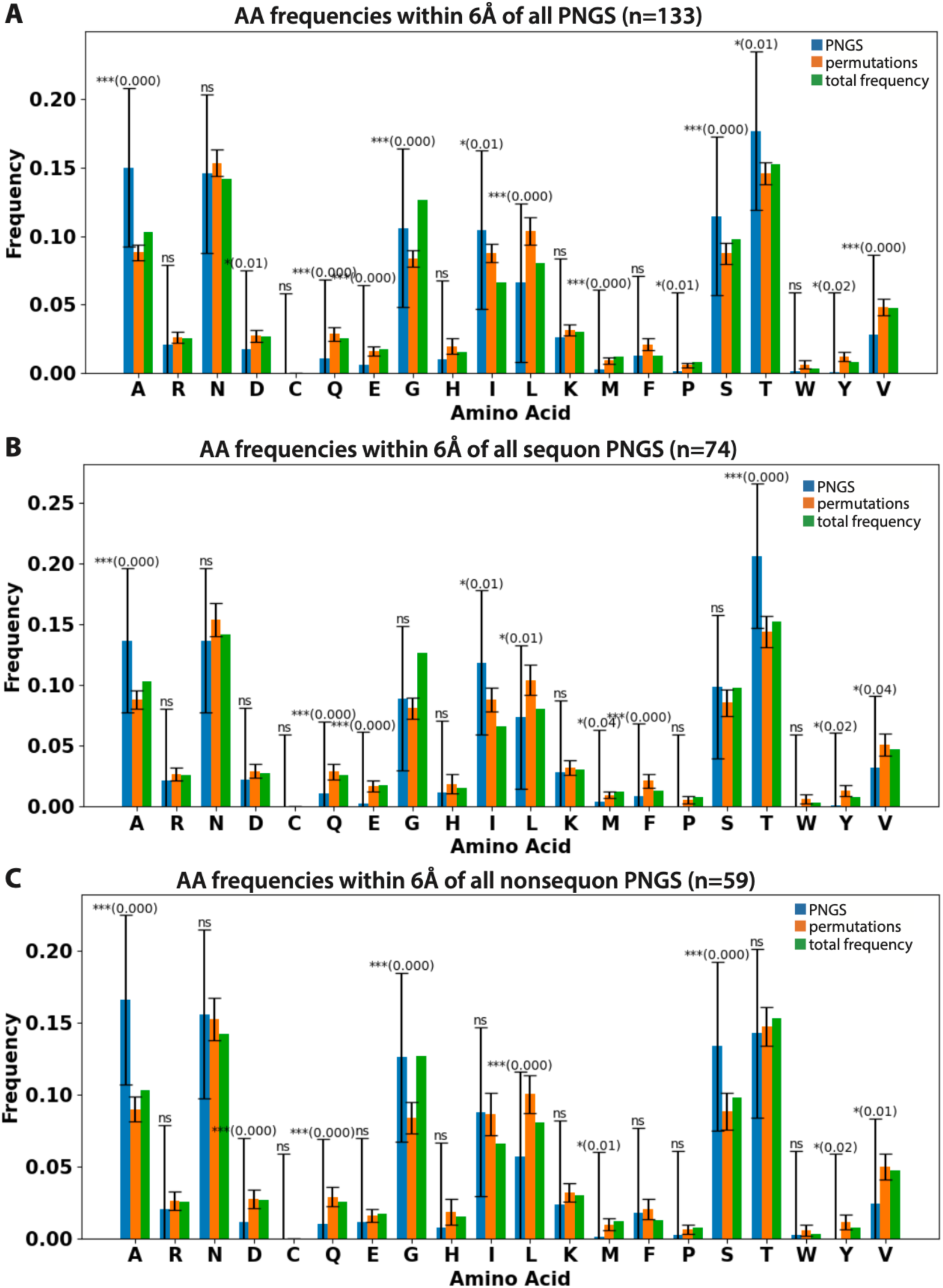
Statistical analysis of local structural environment around PNGS. **A.** Bar plot showing the average frequency of each amino acid type within 6Å of all PNGS, canonical sequon PNGS (**B**), and non-canonical sequon PNGS (**C**). Blue bars are frequencies for the input list of residues, orange bars are the average frequencies across all 1000 permutation samples (with error bars and significance measures as described in Supplemental figure 5), and green bars are the frequency of that amino acid within rEtpA.

**Supplemental figure 7.**
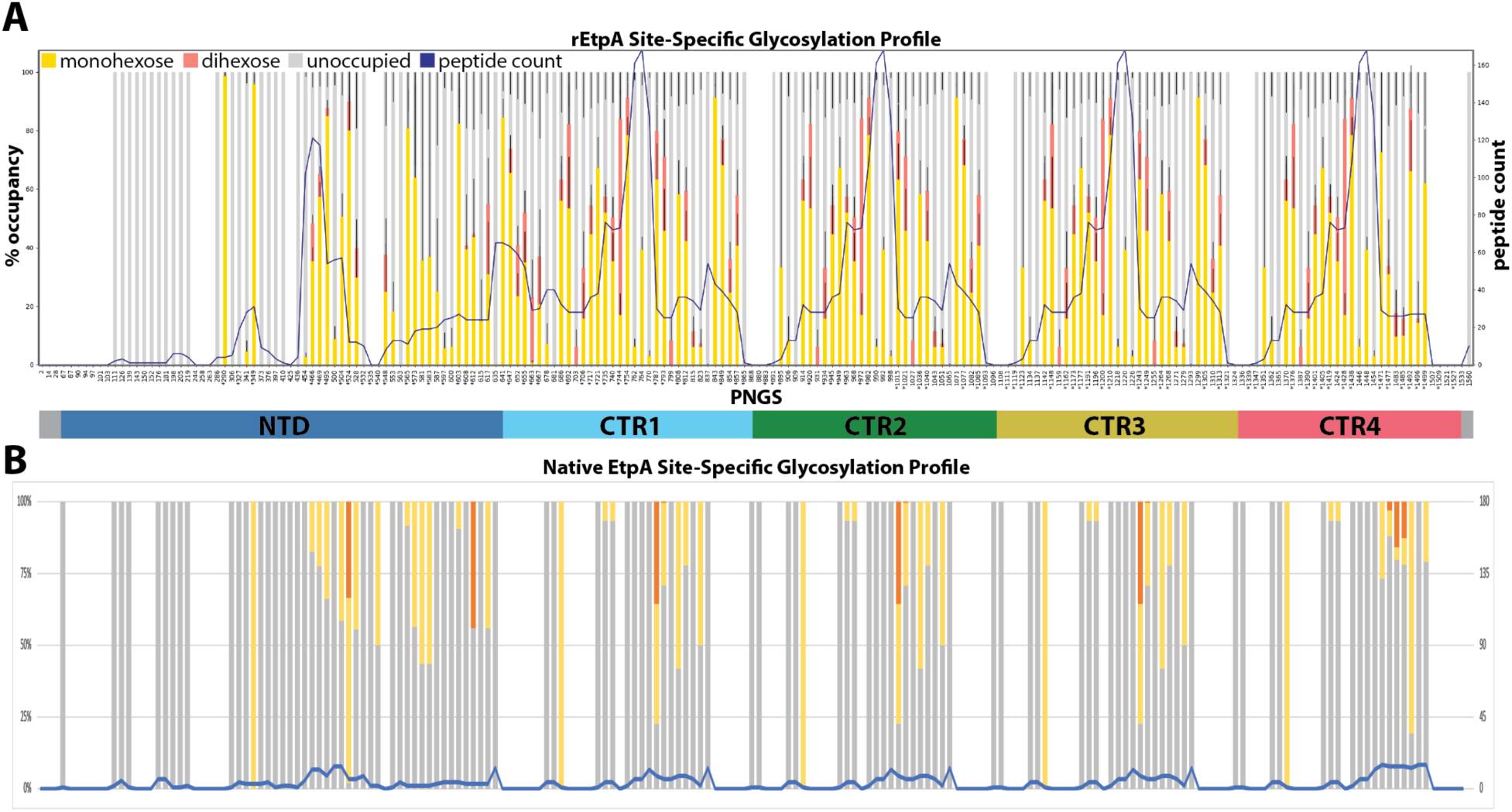
Site-specific glycosylation analysis of native EtpA. **A.** Glycosylation profile for rEtpA reproduced from Figure 3. **B**. Glycosylation profile of native EtpA from ETEC strain H10407. Left axis is % occupancy and right axis is peptide count. Note the much lower peptide counts for native EtpA.

**Supplemental figure 8.**
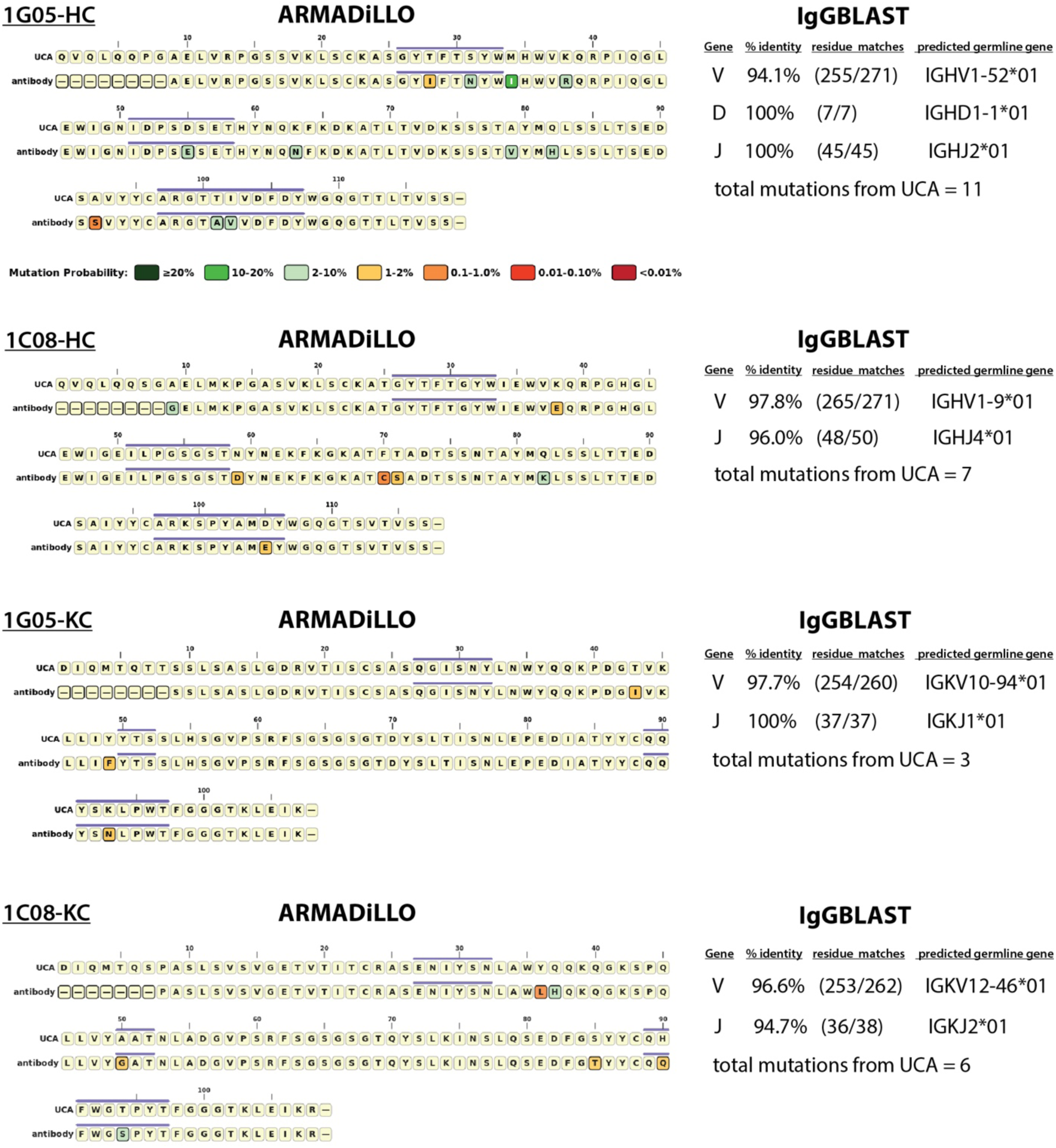
Germline gene IgGBlast and somatic hypermutation analysis by ARMADILLO. IgGBlast results summaries for 1C08 and 1G05 heavy chains (HC) and kappa chains (KC) showing the top match for V, D, and J mouse germline genes. Next to the summary tables are the percent identity to the top matching genes along with the gene names and the total number of somatic hypermutations (SHM) away from the predicted unmutated common ancestor (UCA). Also shown are output plots from the program ARMADILLO with SHM sites shown and scored by their probability, with red being the least probable and green being the most probable.

**Supplemental figure 9.**
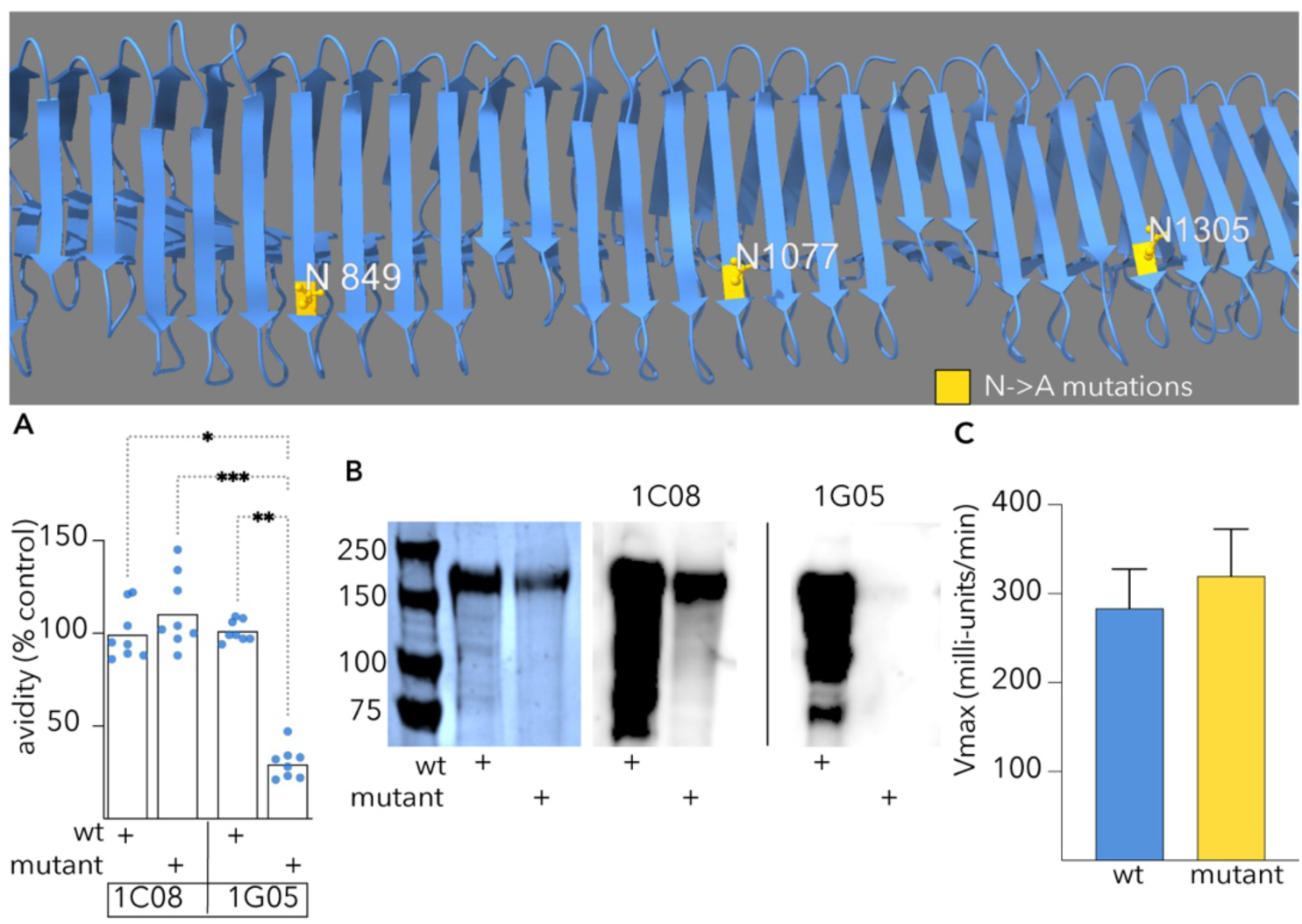
Mutation of 1G05 epitope glycan sites impacts avidity. Figure at top depicts relative location of asparagine (N) to alanine (A) mutations to the putative 1G05 epitope. **A.** mAb 1G05 exhibits decreased avidity for recombinant mutant EtpA with N to A substitutions within C-terminal repeat region at positions 849, 1077, and 1305. Avidity indices (AI) were determined by kinetic ELISA with and without addition of 8 M urea as the chaotropic agent. AI (%) = (Vmax with urea)/(Vmax without urea) and expressed as % of the wild type recombinant protein. Comparisons by Kruskal-Wallis (n=8 technical replicates/group from 2 independent experiments) ***=0.0003, **=0.0037, *=0.03. **B.** Immunoblot recognition of wild type and mutant protein by 1C08 and 1G05. PAGE image (left) indicates protein loading and MW markers. **C.** Blood group A binding by wild type and mutant protein in kinetic ELISA assay.

**Supplemental figure 10.**
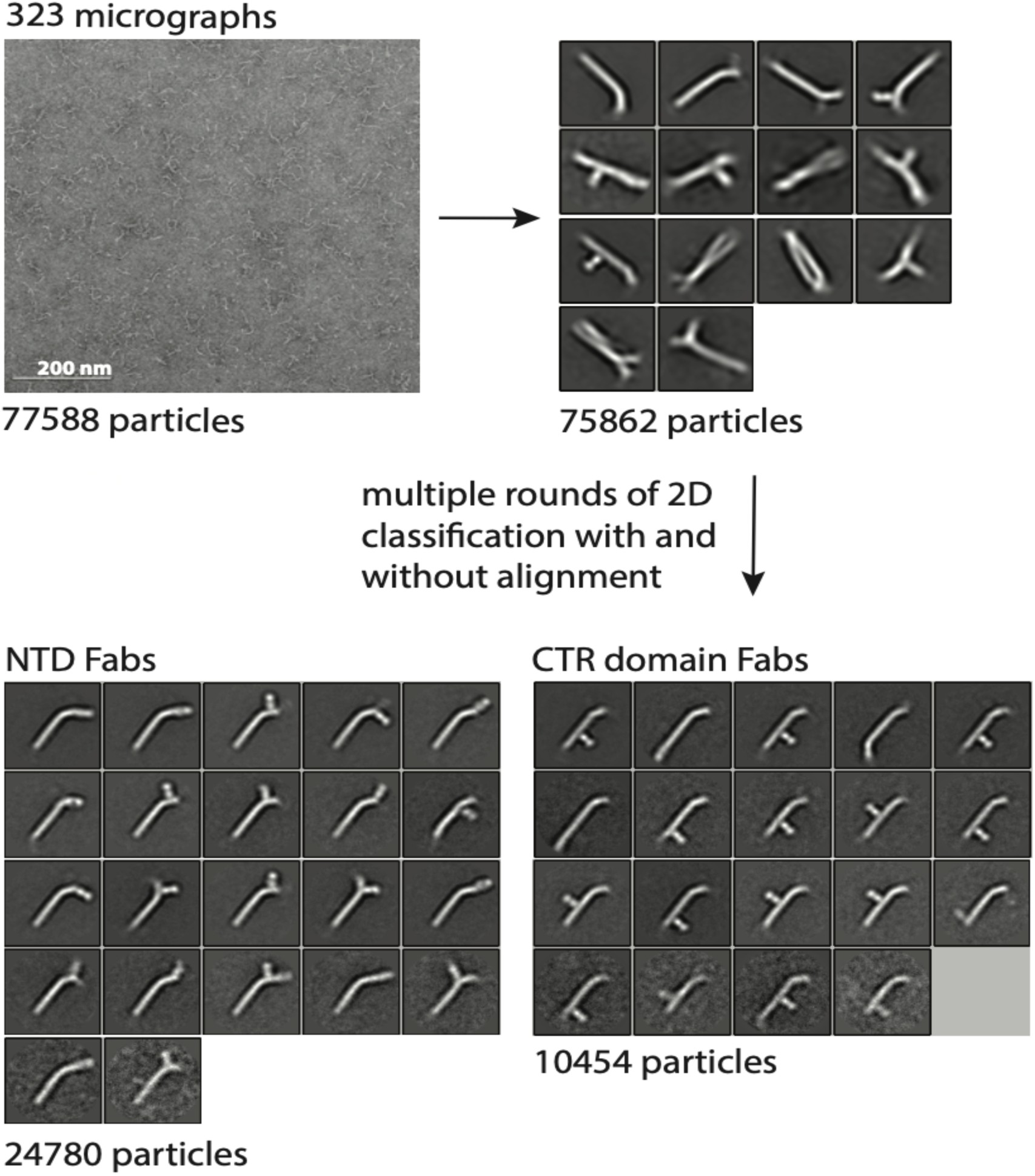
EMPEM processing workflow. Simplified negative stain EMPEM data processing workflow including a representative micrograph, 2D class averages, and particle counts at each step.

**Supplemental figure 11.**
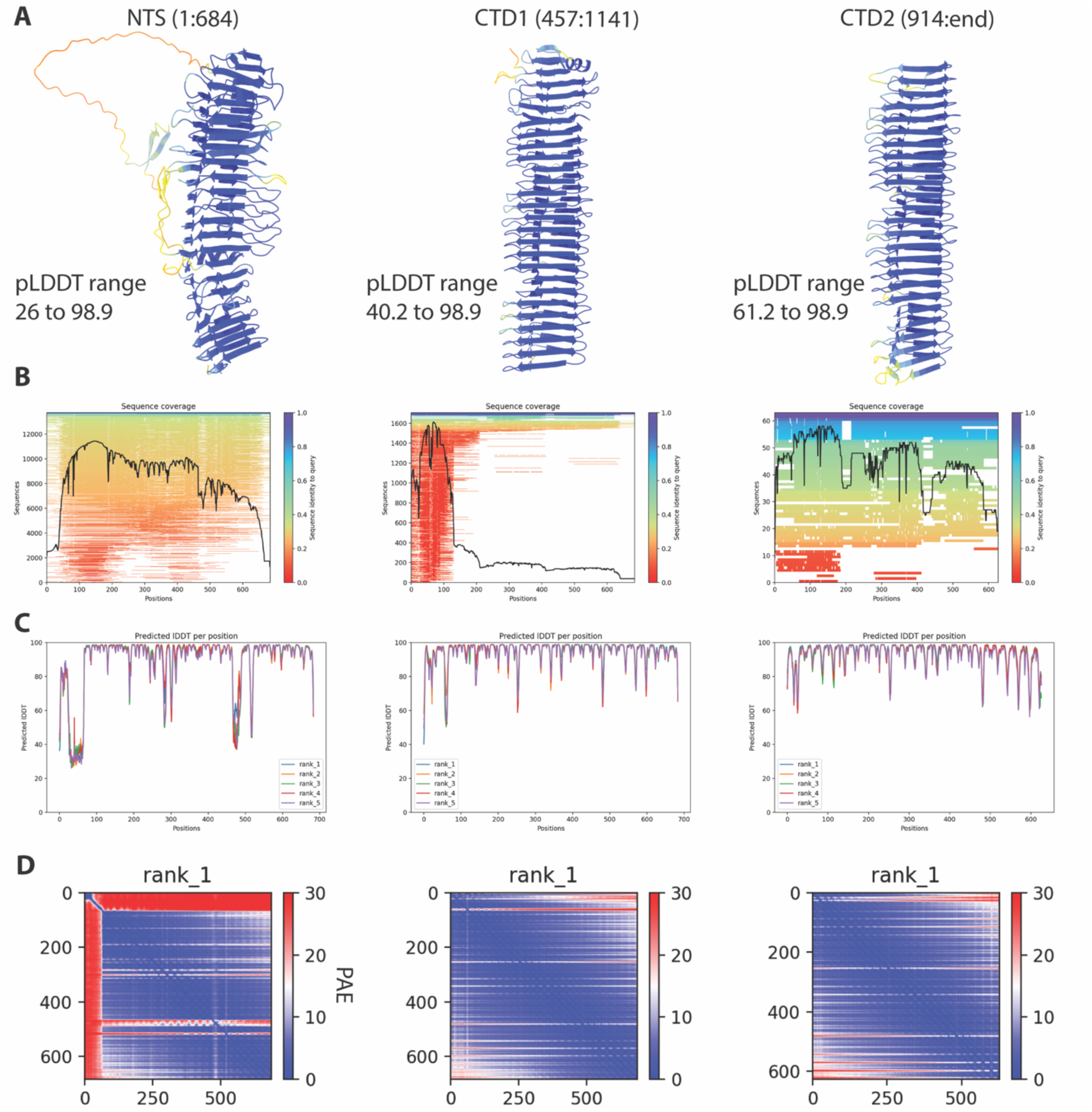
AlphaFold2 predictions for full-length EtpA. **A**. AlphaFold2 top ranking model colored by pLDDT score (blue = high). **B**. Per-residue sequence coverage and identity. **C**. Per-residue pLDDT scores. **D**. PAE (predicted aligned error) matrix.

### Supplemental tables

**Supplemental table 1:**

| supplemental table 1 strains and plasmids |  |  |  |
| --- | --- | --- | --- |
| strain | description/genotype | reference |  |
| Top10 | F-mcrA (mrr-hsdRMS-mcrBC) 80lacZM15 lacX74 recA1 araD139 (ara-leu)7697 galU galK - rpsL(StrR) endA1 nupG | Invitrogen |  |
| jf1696 | Top10(pJL017/pJL030), AmpR, CmR | 48,71 |  |
| jf2826 | LMG194 *fliC*::KmR | 48 |  |
| jf3013 | Top10(pQL211/pJL030), AmpR, CmR | this study |  |
| jf3099 | H10407 *fliC*::KmR | 48 |  |
| jf5090 | Top10(pMH4/pJL030) AmpR, CmR | this study |  |
| jf5381 | jf2826(pJL030) |  |  |
| jf5500 | jf5381(pJL030/pTV005) AmpR, CmR | this study |  |
| H10407 | wild type ETEC strain *etpBAC* serotype O78:H11, LT/STh/STp | 17,78 |  |
| plasmid | description | reference(s) | addgene number |
| pJL030 | pACYC184 based *etpC* expression plasmid CmR | 48,123 | 53533 |
| pJL017 | *etpBA* cloned into pBAD/Myc- His A*, with *etpA* in-frame with myc and 6His coding regions | [35] | 53532 |
| pJMF1028 | linker scanning mutant of pJL017 with transprimer insertion at 3258 of *etpA* | 48 |  |
| pMH4 | EtpA-amplified from pJMF1028, encodes EtpA (AA 1-1086) | this study |  |
| pQL211 | N-terminal region EtpA subclone (AA 1-607) generated with primers jf02214.3/jf022114.4 cloned in-frame with myc and 6His coding regions of pBAD/Myc- His A. | 124 |  |
| pTV005 | Mutant EtpA expression plasmid bearing N849A, N1077A, and N1305A mutations | this study |  |

KmR=kanamycin resistant; AmpR=beta-lactamase/ampicillin resistant; CmR=chloramphenicol resistant.

**Supplemental table 2:**
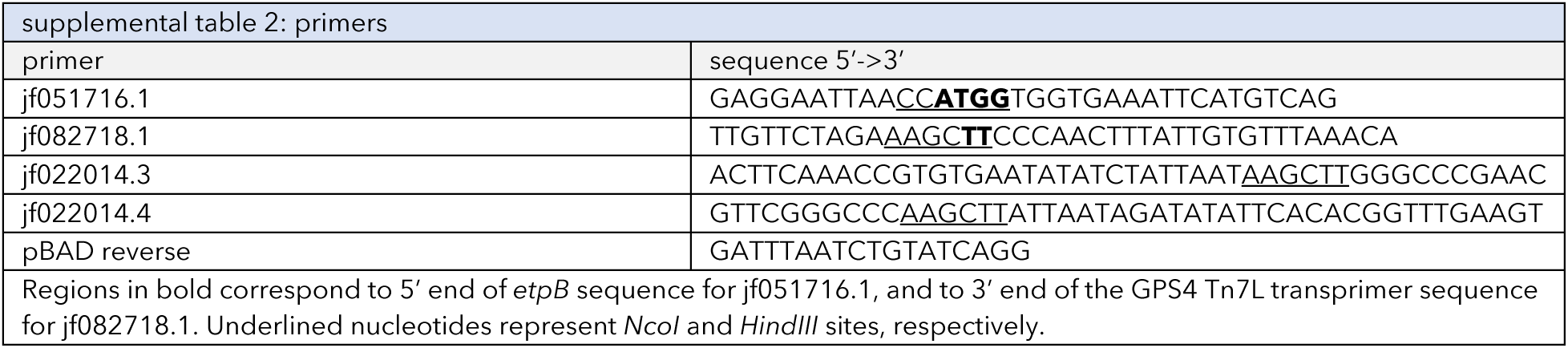

**Supplemental table 3:** Cryo-EM data collection, refinement, and validation statistics

|  | rEtpA:1C08 Fab<br>(EMDB-xxxx)<br>(PDB xxxx) | rEtpA:1G05 Fab<br>(EMDB-xxxx)<br>(PDB xxxx) |
| --- | --- | --- |
| **Data collection and processing** |  |  |
| Magnification | 97,0000 | 97,0000 |
| Voltage (kV) | 200 | 200 |
| Electron exposure (e-/Å<sup>2</sup>) | 47 | 47 |
| Defocus range (μm) | ~-0.25:-2.0um | ~-0.4:-2.0um |
| Pixel size (Å) | 0.725Å | 0.725Å |
| Symmetry imposed | none | none |
| Initial particle images (no.) | 4,178,645 | 1,651,766 |
| Final particle images (no.) | 215,360 | 52,107 |
| Map resolution (Å) | 3.34Å | 3.97Å |
| FSC threshold | 0.143 | 0.143 |
| Map resolution range (Å) | ~3-5Å | ~3.5-5Å |
| Map sharpening B factor (Å<sup>2</sup>) | 110.5 | 380.4 |
| **Refinement** |  |  |
| **Initial model used (PDB code)** | **AlphaFold2 and**<br>**SABPred**<br>**predictions** | **AlphaFold2 and**<br>**SABPred**<br>**predictions** |
| <u>Model composition</u> |  |  |
| Non-hydrogen atoms | 9416 | 9444 |
| Protein residues | 1260 | 1260 |
| Ligands | 39 BGC | 38 BGC |
| <u>R.m.s. deviations</u> |  |  |
| Bond lengths (Å) | 0.011 | 0.01 |
| Bond angles (°) | 1.318 | 1.03 |
| <u>Validation</u> |  |  |
| MolProbity score | 2.03 | 2.02 |
| Clashscore | 14.87 | 15.05 |
| Poor rotamers (%) | 0.51 | 0.41 |
| CaBLAM outliers (%) | 2.8 | 3.53 |
| <u>Ramachandran plot</u> |  |  |
| Favored (%) | 94.90 | 95.14 |
| Allowed (%) | 4.63 | 4.63 |
| Disallowed (%) | 0.48 | 0.24 |
| EMRinger score | 3.6 | 2.7 |

## References

1. Fleckenstein, J.M., and Sheikh, A. (2021). Emerging Themes in the Molecular Pathogenesis of Enterotoxigenic Escherichia coli. The Journal of infectious diseases 224, S813--S820.

2. Khalil, I.A., Troeger, C., Blacker, B.F., Rao, P.C., Brown, A., Atherly, D.E., Brewer, T.G., Engmann, C.M., Houpt, E.R., Kang, G., and, et al. (2018). Morbidity and mortality due to shigella and enterotoxigenic Escherichia coli diarrhoea: the Global Burden of Disease Study 1990--2016. The Lancet infectious diseases 18, 1229--1240.

3. Kotloff, K.L., Nataro, J.P., Blackwelder, W.C., Nasrin, D., Farag, T.H., Panchalingam, S., Wu, Y., Sow, S.O., Sur, D., Breiman, R.F., and, et al. (2013). Burden and aetiology of diarrhoeal disease in infants and young children in developing countries (the Global Enteric Multicenter Study, GEMS): a prospective, case-control study. The Lancet 382, 209--222.

4. Levine, M.M., Nasrin, D., Accio, S., Bassat, Q., Powell, H., Tennant, S.M., Sow, S.O., Sur, D., Zaidi, A.K.M., Faruque, A.S.G., and, et al. (2020). Diarrhoeal disease and subsequent risk of death in infants and children residing in low-income and middle-income countries: analysis of the GEMS case-control study and 12-month GEMS-1A follow-on study. The Lancet Global Health 8, e204--e214.

5. Nasrin, D., Blackwelder, W.C., Sommerfelt, H., Wu, Y., Farag, T.H., Panchalingam, S., Biswas, K., Saha, D., Jahangir Hossain, M., Sow, S.O., and, et al. (2021). Pathogens associated with linear growth faltering in children with diarrhea and impact of antibiotic treatment: the global enteric multicenter study. The Journal of infectious diseases 224, S848--S855.

6. Mapping child growth failure across low-and middle-income countries. (2020). Nature 577, 231--234.

7. Black, R.E., Brown, K.H., and Becker, S. (1984). Effects of diarrhea associated with specific enteropathogens on the growth of children in rural Bangladesh. Pediatrics 73, 799--805.

8. Anderson Iv, J.D., Bagamian, K.H., Muhib, F., Amaya, M.P., Laytner, L.A., Wierzba, T., and Rheingans, R. (2019). Burden of enterotoxigenic Escherichia coli and shigella non-fatal diarrhoeal infections in 79 low-income and lower middle-income countries: a modelling analysis. The Lancet Global Health 7, e321--e330.

9. Qadri, F., Saha, A., Ahmed, T., Al Tarique, A., Begum, Y.A., and Svennerholm, A.-M. (2007). Disease burden due to enterotoxigenic Escherichia coli in the first 2 years of life in an urban community in Bangladesh. Infection and immunity 75, 3961--3968.

10. Platts-Mills, J.A., Taniuchi, M., Uddin, M.J., Sobuz, S.U., Mahfuz, M., Gaffar, S.M.A., Mondal, D., Hossain, M.I., Islam, M.M., Ahmed, A.M.S., and, et al. (2017). Association between enteropathogens and malnutrition in children aged 6--23 mo in Bangladesh: a case-control study. The American journal of clinical nutrition 105, 1132--1138.

11. Mondal, D., Haque, R., Sack, R.B., Kirkpatrick, B.D., and Petri Jr, W.A. (2009). AeribuCon of malnutrition to cause-specific diarrheal illness: evidence from a prospective study of preschool children in Mirpur, Dhaka, Bangladesh. The American Journal of Tropical Medicine and Hygiene 80, 824.

12. Mondal, D., Minak, J., Alam, M., Liu, Y., Dai, J., Korpe, P., Liu, L., Haque, R., and Petri Jr, W.A. (2012). Contribution of enteric infection, altered intestinal barrier function, and maternal malnutrition to infant malnutrition in Bangladesh. Clinical Infectious Diseases 54, 185--192.

13. Kotloff, K.L., Nasrin, D., Blackwelder, W.C., Wu, Y., Farag, T., Panchalingham, S., Sow, S.O., Sur, D., Zaidi, A.K.M., Faruque, A.S.G., and, et al. (2019). The incidence, aetiology, and adverse clinical consequences of less severe diarrhoeal episodes among infants and children residing in low-income and middle-income countries: a 12-month case-control study as a follow-on to the Global Enteric Multicenter Study (GEMS). The Lancet Global Health 7, e568--e584.

14. Organization, W.H. (2021). WHO preferred product characteristics for vaccines against enterotoxigenic Escherichia coli. https://www.who.int/publicaCons/i/item/who-preferred-product-characterisCcs-for-vaccines-against-enterotoxigenic-escherichia-coli.

15. Hosangadi, D., Smith, P.G., and Giersing, B.K. (2019). Considerations for using ETEC and Shigella disease burden estimates to guide vaccine development strategy. Vaccine 37, 7372--7380.

16. Khalil, I., Anderson, J.D., Bagamian, K.H., Baqar, S., Giersing, B., Hausdorff, W.P., Marshall, C., Porter, C.K., Walker, R.I., and Bourgeois, A.L. (2023). Vaccine value profile for enterotoxigenic Escherichia coli (ETEC). Vaccine 41 Suppl 2, S95–S113. 10.1016/j.vaccine.2023.02.011.

17. Fleckenstein, J.M., Roy, K., Fischer, J.F., and Burkitt, M. (2006). Identification of a two-partner secretion locus of enterotoxigenic Escherichia coli. Infection and immunity 74, 2245–2258.

18. Clantin, B., Delattre, A.-S., Rucktooa, P., Saint, N., Mli, A.C., Locht, C., Jacob-Dubuisson, F., and Villeret, V. (2007). Structure of the membrane protein FhaC: a member of the Omp85-TpsB transporter superfamily. Science 317, 957--961.

19. Roy, K., Hilliard, G.M., Hamilton, D.J., Luo, J., Ostmann, M.M., and Fleckenstein, J.M. (2009). Enterotoxigenic Escherichia coli EtpA mediates adhesion between flagella and host cells. Nature 457, 594--598.

20. Dorsey, F.C., Fischer, J.F., and Fleckenstein, J.M. (2006). Directed delivery of heat-labile enterotoxin by enterotoxigenic Escherichia coli. Cellular microbiology 8, 1516--1527.

21. Roy, K., Hamilton, D.J., and Fleckenstein, J.M. (2012). Cooperative role of antibodies against heat-labile toxin and the EtpA adhesin in preventing toxin delivery and intestinal colonization by enterotoxigenic Escherichia coli. Clinical and Vaccine Immunology 19, 1603--1608.

22. Zhu, Y., Luo, Q., Davis, S.M., Westra, C., Vickers, T.J., and Fleckenstein, J.M. (2018). Molecular determinants of enterotoxigenic Escherichia coli heat-stable toxin secretion and delivery. Infection and immunity 86, e00526--00518.

23. Kumar, P., Kuhlmann, F.M., Bhullar, K., Yang, H., Vallance, B.A., Xia, L., Luo, Q., and Fleckenstein, J.M. (2016). Dynamic Interactions of a Conserved Enterotoxigenic Escherichia coli Adhesin with Intestinal Mucins Govern Epithelium Engagement and Toxin Delivery. Infection and immunity 84, 3608–3617. 10.1128/IAI.00692-16.

24. Kumar, P., Kuhlmann, F.M., Chakraborty, S., Bourgeois, A.L., Foulke-Abel, J., Tumala, B., Vickers, T.J., Sack, D.A., DeNearing, B., Harro, C.D., and, et al. (2018). Enterotoxigenic Escherichia coli--blood group A interactions intensify diarrheal severity. The Journal of clinical investigation 128, 3298--3311.

25. Sahl, J.W., Steinsland, H., Redman, J.C., Angiuoli, S.V., Nataro, J.P., Sommerfelt, H., and Rasko, D.A. (2011). A comparative genomic analysis of diverse clonal types of enterotoxigenic Escherichia coli reveals pathovar-specific conservation. Infection and immunity 79, 950--960.

26. Kuhlmann, F.M., Martin, J., Hazen, T.H., Vickers, T.J., Pashos, M., Okhuysen, P.C., Gmez-Duarte, O.G., Cebelinski, E., Boxrud, D., Del Canto, F., and, et al. (2019). Conservation and global distribution of non-canonical antigens in Enterotoxigenic Escherichia coli. PLoS neglected tropical diseases 13, e0007825.

27. Luo, Q., Qadri, F., Kansal, R., Rasko, D.A., Sheikh, A., and Fleckenstein, J.M. (2015). Conservation and immunogenicity of novel antigens in diverse isolates of enterotoxigenic Escherichia coli. PLoS neglected tropical diseases 9, e0003446.

28. Mondal, I., Bhakat, D., Chowdhury, G., Manna, A., Samanta, S., Deb, A.K., Mukhopadhyay, A.K., and Chatterjee, N.S. (2022). Distribution of virulence factors and its relatedness towards the antimicrobial response of enterotoxigenic Escherichia coli strains isolated from patients in Kolkata, India. Journal of Applied Microbiology 132, 675--686.

29. von Mentzer, A., Connor, T.R., Wieler, L.H., Semmler, T., Iguchi, A., Thomson, N.R., Rasko, D.A., Joffre, E., Corander, J., Pickard, D., and, et al. (2014). Identification of enterotoxigenic Escherichia coli (ETEC) clades with long-term global distribution. Nature genetics 46, 1321--1326.

30. Kuhlmann, F.M., Laine, R.O., Afrin, S., Nakajima, R., Akhtar, M., Vickers, T., Parker, K., Nizam, N.N., Grigura, V., Goss, C.W., and, et al. (2021). Contribution of noncanonical antigens to virulence and adaptive immunity in human infection with enterotoxigenic E. coli. Infection and immunity 89, e00041--00021.

31. Chakraborty, S., Randall, A., Vickers, T.J., Molina, D., Harro, C.D., DeNearing, B., Brubaker, J., Sack, D.A., Bourgeois, A.L., Felgner, P.L., and, et al. (2019). Interrogation of a live-attenuated enterotoxigenic Escherichia coli vaccine highlights features unique to wild-type infection. NPJ vaccines 4, 1--9.

32. Chakraborty, S., Randall, A., Vickers, T.J., Molina, D., Harro, C.D., DeNearing, B., Brubaker, J., Sack, D.A., Bourgeois, A.L., Felgner, P.L., and, et al. (2018). Human experimental challenge with enterotoxigenic Escherichia coli elicits immune responses to canonical and novel antigens relevant to vaccine development. The Journal of infectious diseases 218, 1436--1446.

33. von Mentzer, A., Blackwell, G.A., Pickard, D., Boinett, C.J., and Joffr (2021). Long-read-sequenced reference genomes of the seven major lineages of enterotoxigenic Escherichia coli (ETEC) circulating in modern Cme. Scientific reports 11, 1--16.

34. Luo, Q., Vickers, T.J., and Fleckenstein, J.M. (2016). Immunogenicity and protective efficacy against enterotoxigenic Escherichia coli colonization following intradermal, sublingual, or oral vaccination with EtpA adhesin. Clinical and Vaccine Immunology 23, 628--637.

35. Roy, K., Hamilton, D.J., Munson, G.P., and Fleckenstein, J.M. (2011). Outer membrane vesicles induce immune responses to virulence proteins and protect against colonization by enterotoxigenic Escherichia coli. Clinical and vaccine immunology 18, 1803--1808.

36. Roy, K., Hamilton, D., Ostmann, M.M., and Fleckenstein, J.M. (2009). Vaccination with EtpA glycoprotein or flagellin protects against colonization with enterotoxigenic Escherichia coli in a murine model. Vaccine 27, 4601--4608.

37. Roy, K., Hamilton, D., Allen, K.P., Randolph, M.P., and Fleckenstein, J.M. (2008). The EtpA exoprotein of enterotoxigenic Escherichia coli promotes intestinal colonization and is a protective antigen in an experimental model of murine infection. Infection and immunity 76, 2106--2112.

38. Fleckenstein, J., Sheikh, A., and Qadri, F. (2014). Novel antigens for enterotoxigenic Escherichia coli vaccines. Expert review of vaccines 13, 631--639.

39. Walker, R., Kaminski, R.W., Porter, C., Choy, R.K.M., White, J.A., Fleckenstein, J.M., Cassels, F., and Bourgeois, L. (2021). Vaccines for protecting infants from bacterial causes of diarrheal disease. Microorganisms 9, 1382.

40. Khalil, I., Walker, R., Porter, C.K., Muhib, F., Chilengi, R., Cravioto, A., Guerrant, R., Svennerholm, A.-M., Qadri, F., Baqar, S., and, et al. (2021). Enterotoxigenic Escherichia coli (ETEC) vaccines: Priority activities to enable product development, licensure, and global access. Vaccine 39, 4266--4277.

41. Clantin, B., Hodak, H., Willery, E., Locht, C., Jacob-Dubuisson, F., and Villeret, V. (2004). The crystal structure of filamentous hemagglutinin secretion domain and its implications for the two-partner secretion pathway. Proc Natl Acad Sci U S A 101, 6194–6199. 10.1073/pnas.0400291101.

42. Baelen, S., Dewitte, F., Clantin, B., and Villeret, V. (2013). Structure of the secretion domain of HxuA from Haemophilus influenzae. Acta Crystallogr Sect F Struct Biol Cryst Commun 69, 1322–1327. 10.1107/S174430911302962X.

43. Yeo, H.J., Yokoyama, T., Walkiewicz, K., Kim, Y., Grass, S., and Geme, J.W., 3rd (2007). The structure of the Haemophilus influenzae HMW1 pro-piece reveals a structural domain essential for bacterial two-partner secretion. The Journal of biological chemistry 282, 31076–31084. 10.1074/jbc.M705750200.

44. Weaver, T.M., Hocking, J.M., Bailey, L.J., Wawrzyn, G.T., Howard, D.R., Sikkink, L.A., Ramirez-Alvarado, M., and Thompson, J.R. (2009). Structural and functional studies of truncated hemolysin A from Proteus mirabilis. The Journal of biological chemistry 284, 22297–22309. 10.1074/jbc.M109.014431.

45. Morse, R.P., Nikolakakis, K.C., Willett, J.L., Gerrick, E., Low, D.A., Hayes, C.S., and Goulding, C.W. (2012). Structural basis of toxicity and immunity in contact-dependent growth inhibition (CDI) systems. Proc Natl Acad Sci U S A 109, 21480–21485. 10.1073/pnas.1216238110.

46. Zambolin, S., Clantin, B., Chami, M., Hoos, S., Haouz, A., Villeret, V., and Delepelaire, P. (2016). Structural basis for haem piracy from host haemopexin by Haemophilus influenzae. Nature communications 7, 11590. 10.1038/ncomms11590.

47. Guerin, J., Bigot, S., Schneider, R., Buchanan, S.K., and Jacob-Dubuisson, F. (2017). Two-Partner Secretion: Combining Efficiency and Simplicity in the Secretion of Large Proteins for Bacteria-Host and Bacteria-Bacteria Interactions. Front Cell Infect Microbiol 7, 148. 10.3389/fcimb.2017.00148.

48. Roy, K., Hilliard, G.M., Hamilton, D.J., Luo, J., Ostmann, M.M., and Fleckenstein, J.M. (2009). Enterotoxigenic Escherichia coli EtpA mediates adhesion between flagella and host cells. Nature 457, 594–598. 10.1038/nature07568.

49. Nash, Z.M., and Cotter, P.A. (2019). Bordetella Filamentous Hemagglutinin, a Model for the Two-Partner Secretion Pathway. Microbiol Spectr 7. 10.1128/microbiolspec.PSIB-0024-2018.

50. Relman, D.A., Domenighini, M., Tuomanen, E., Rappuoli, R., and Falkow, S. (1989). Filamentous hemagglutinin of Bordetella pertussis: nucleotide sequence and crucial role in adherence. Proc Natl Acad Sci U S A 86, 2637–2641.

51. Lis, H., and Sharon, N. (1998). Lectins: carbohydrate-specific proteins that mediate cellular recognition. Chemical reviews 98, 637--674.

52. Mahajan, S., and Ramya, T.N.C. (2018). Nature-inspired engineering of an F-type lectin for increased binding strength. Glycobiology 28, 933--948.

53. Elgavish, S., and Shaanan, B. (1997). Lectin-carbohydrate interactions: different folds, common recognition principles. Trends in biochemical sciences 22, 462--467.

54. Notova, S., Bonnardel, F., Lisacek, F., Varrot, A., and Imberty, A. (2020). Structure and engineering of tandem repeat lectins. Current Opinion in Structural Biology 62, 39--47.

55. Grass, S., and St Geme, J.W., 3rd (2000). Maturation and secretion of the non-typable Haemophilus influenzae HMW1 adhesin: roles of the N-terminal and C-terminal domains. Molecular microbiology 36, 55–67.

56. Herron, S.R., Benen, J.A., Scavetta, R.D., Visser, J., and Jurnak, F. (2000). Structure and function of pectic enzymes: virulence factors of plant pathogens. Proc Natl Acad Sci U S A 97, 8762–8769. 10.1073/pnas.97.16.8762.

57. Ntui, C.M., Fleckenstein, J.M., and Schubert, W.D. (2023). Structural and biophysical characterization of the secreted, beta-helical adhesin EtpA of Enterotoxigenic Escherichia coli. PLoS One 18, e0287100. 10.1371/journal.pone.0287100.

58. Grass, S., Buscher, A.Z., Swords, W.E., Apicella, M.A., Barenkamp, S.J., Ozchlewski, N., and St Geme Iii, J.W. (2003). The Haemophilus influenzae HMW1 adhesin is glycosylated in a process that requires HMW1C and phosphoglucomutase, an enzyme involved in lipooligosaccharide biosynthesis. Molecular microbiology 48, 737--751.

59. Grass, S., Lichti, C.F., Townsend, R.R., Gross, J., and St. Geme Iii, J.W. (2010). The Haemophilus influenzae HMW1C protein is a glycosyltransferase that transfers hexose residues to asparagine sites in the HMW1 adhesin. PLoS pathogens 6, e1000919.

60. McCann, J.R., and St. Geme Iii, J.W. (2014). The HMW1C-like glycosyltransferases—an enzyme family with a sweet tooth for simple sugars. PLoS pathogens 10, e1003977.

61. Jumper, J., Evans, R., Pritzel, A., Green, T., Figurnov, M., Ronneberger, O., Tunyasuvunakool, K., Bates, R., Zidek, A., Potapenko, A., et al. (2021). Highly accurate protein structure prediction with AlphaFold. Nature 596, 583–589. 10.1038/s41586-021-03819-2.

62. Kawai, F., Grass, S., Kim, Y., Choi, K.J., St Geme, J.W., 3rd, and Yeo, H.J. (2011). Structural insights into the glycosyltransferase activity of the Actinobacillus pleuropneumoniae HMW1C-like protein. J Biol Chem 286, 38546–38557. 10.1074/jbc.M111.237602.

63. Gross, J., Grass, S., Davis, A.E., Gilmore-Erdmann, P., Townsend, R.R., and Geme, J.W.S. (2008). The Haemophilus influenzae HMW1 adhesin is a glycoprotein with an unusual N-linked carbohydrate modification. Journal of Biological Chemistry 283, 26010--26015.

64. Fleckenstein, J.M., Roy, K., Fischer, J.F., and Burkitt, M. (2006). Identification of a two-partner secretion locus of enterotoxigenic Escherichia coli. Infection and immunity 74, 2245--2258.

65. Baboo, S., Diedrich, J.K., Martinez-Bartolome, S., Wang, X., Schiffner, T., Groschel, B., Schief, W.R., Paulson, J.C., and Yates, J.R., 3rd (2023). DeGlyPHER: Highly sensitive site-specific analysis of N-linked glycans on proteins. Methods Enzymol 682, 137–185. 10.1016/bs.mie.2022.09.004.

66. Gross, J., Grass, S., Davis, A.E., Gilmore-Erdmann, P., Townsend, R.R., and St Geme, J.W., 3rd (2008). The Haemophilus influenzae HMW1 adhesin is a glycoprotein with an unusual N-linked carbohydrate modification. The Journal of biological chemistry 283, 26010–26015. 10.1074/jbc.M801819200.

67. Watanabe, Y., Allen, J.D., Wrapp, D., McLellan, J.S., and Crispin, M. (2020). Site-specific glycan analysis of the SARS-CoV-2 spike. Science 369, 330–333. 10.1126/science.abb9983.

68. Ardejani, M.S., Noodleman, L., Powers, E.T., and Kelly, J.W. (2021). Stereoelectronic effects in stabilizing protein-N-glycan interactions revealed by experiment and machine learning. Nat Chem 13, 480–487. 10.1038/s41557-021-00646-w.

69. Barb, A.W., Borgert, A.J., Liu, M., Barany, G., and Live, D. (2010). Chapter Eighteen - Intramolecular Glycan–Protein Interactions in Glycoproteins. In Methods in Enzymology, M. Fukuda, ed. (Academic Press), pp. 365–388. 10.1016/S0076-6879(10)78018-6.

70. Martin Beem, J.S., Venkatayogi, S., Haynes, B.F., and Wiehe, K. (2023). ARMADiLLO: a web server for analyzing antibody mutation probabilities. Nucleic Acids Res 51, W51–w56. 10.1093/nar/gkad398.

71. Roy, K., Hamilton, D., Allen, K.P., Randolph, M.P., and Fleckenstein, J.M. (2008). The EtpA exoprotein of enterotoxigenic Escherichia coli promotes intestinal colonization and is a protective antigen in an experimental model of murine infection. Infection and immunity 76, 2106–2112. IAI.01304-07 [pii] 10.1128/IAI.01304-07.

72. Roy, K., Hamilton, D., Ostmann, M.M., and Fleckenstein, J.M. (2009). Vaccination with EtpA glycoprotein or flagellin protects against colonization with enterotoxigenic Escherichia coli in a murine model. Vaccine 27, 4601–4608. 10.1016/j.vaccine.2009.05.076.

73. Roy, K., Hamilton, D.J., and Fleckenstein, J.M. (2012). Cooperative role of antibodies against heat-labile toxin and the EtpA Adhesin in preventing toxin delivery and intestinal colonization by enterotoxigenic Escherichia coli. Clinical and vaccine immunology : CVI 19, 1603–1608. 10.1128/CVI.00351-12.

74. von Mentzer, A., and Svennerholm, A.M. (2023). Colonization factors of human and animal-specific enterotoxigenic Escherichia coli (ETEC). Trends Microbiol. 10.1016/j.Cm.2023.11.001.

75. Kuhlmann, F.M., Martin, J., Hazen, T.H., Vickers, T.J., Pashos, M., Okhuysen, P.C., Gomez-Duarte, O.G., Cebelinski, E., Boxrud, D., Del Canto, F., et al. (2019). Conservation and global distribution of non-canonical antigens in Enterotoxigenic Escherichia coli. PLoS Negl Trop Dis 13, e0007825. 10.1371/journal.pntd.0007825.

76. Sack, R.B., Gorbach, S.L., Banwell, J.G., Jacobs, B., Chatterjee, B.D., and Mitra, R.C. (1971). Enterotoxigenic Escherichia coli isolated from patients with severe cholera-like disease. J Infect Dis 123, 378–385.

77. Dorsey, F.C., Fischer, J.F., and Fleckenstein, J.M. (2006). Directed delivery of heat-labile enterotoxin by enterotoxigenic Escherichia coli. Cell Microbiol 8, 1516–1527. 10.1111/j.1462-5822.2006.00736.x.

78. Evans, D.J., Jr., and Evans, D.G. (1973). Three characteristics associated with enterotoxigenic Escherichia coli isolated from man. Infection and immunity 8, 322–328. DOI: 10.1128/iai.8.3.322-328.1973.

79. Kumar, P., Kuhlmann, F.M., Chakraborty, S., Bourgeois, A.L., Foulke-Abel, J., Tumala, B., Vickers, T.J., Sack, D.A., DeNearing, B., Harro, C.D., et al. (2018). Enterotoxigenic Escherichia coli-blood group A interactions intensify diarrheal severity. The Journal of clinical investigation 128, 3298–3311. 10.1172/JCI97659.

80. Qadri, F., Saha, A., Ahmed, T., Al Tarique, A., Begum, Y.A., and Svennerholm, A.M. (2007). Disease burden due to enterotoxigenic Escherichia coli in the first 2 years of life in an urban community in Bangladesh. Infection and immunity 75, 3961–3968. 10.1128/IAI.00459-07.

81. Boraston, A.B., Wang, D., and Burke, R.D. (2006). Blood group antigen recognition by a Streptococcus pneumoniae virulence factor. The Journal of biological chemistry 281, 35263–35271. 10.1074/jbc.M607620200.

82. Ficko-Blean, E., and Boraston, A.B. (2009). N-acetylglucosamine recognition by a family 32 carbohydrate-binding module from Clostridium perfringens NagH. J Mol Biol 390, 208–220. 10.1016/j.jmb.2009.04.066.

83. van Bueren, A.L., Higgins, M., Wang, D., Burke, R.D., and Boraston, A.B. (2007). Identification and structural basis of binding to host lung glycogen by streptococcal virulence factors. Nature structural & molecular biology 14, 76–84. 10.1038/nsmb1187.

84. Cao, L., Pauthner, M., Andrabi, R., Rantalainen, K., Berndsen, Z., Diedrich, J.K., Menis, S., Sok, D., Bastidas, R., Park, S.R., et al. (2018). Differential processing of HIV envelope glycans on the virus and soluble recombinant trimer. Nat Commun 9, 3693. 10.1038/s41467-018-06121-4.

85. Alam, S.M., Aussedat, B., Vohra, Y., Meyerhoff, R.R., Cale, E.M., Walkowicz, W.E., Radakovich, N.A., Anasti, K., Armand, L., Parks, R., et al. (2017). Mimicry of an HIV broadly neutralizing antibody epitope with a synthetic glycopeptide. Science translational medicine 9. 10.1126/scitranslmed.aai7521.

86. Watanabe, Y., Bowden, T.A., Wilson, I.A., and Crispin, M. (2019). Exploitation of glycosylation in enveloped virus pathobiology. Biochim Biophys Acta Gen Subj 1863, 1480–1497. 10.1016/j.bbagen.2019.05.012.

87. Fleckenstein, J.M., and Roy, K. (2009). Purification of recombinant high molecular weight two-partner secretion proteins from Escherichia coli. Nature protocols 4, 1083--1092.

88. Gutierrez, R.L., Porter, C.K., Harro, C., Talaat, K., Riddle, M.S., DeNearing, B., Brubaker, J., Maciel, M., Jr., Laird, R.M., Poole, S., et al. (2024). Efficacy Evaluation of an Intradermally Delivered Enterotoxigenic Escherichia coli CF Antigen I Fimbrial Tip Adhesin Vaccine Coadministered with Heat-Labile Enterotoxin with LT(R192G) against Experimental Challenge with Enterotoxigenic E. coli H10407 in Healthy Adult Volunteers. Microorganisms 12. 10.3390/microorganisms12020288.

89. von Boehmer, L., Liu, C., Ackerman, S., Gitlin, A.D., Wang, Q., Gazumyan, A., and Nussenzweig, M.C. (2016). Sequencing and cloning of antigen-specific antibodies from mouse memory B cells. Nat Protoc 11, 1908–1923. 10.1038/nprot.2016.102.

90. Alsoussi, W.B., Turner, J.S., Case, J.B., Zhao, H., Schmitz, A.J., Zhou, J.Q., Chen, R.E., Lei, T., Rizk, A.A., McIntire, K.M., et al. (2020). A Potently Neutralizing Antibody Protects Mice against SARS-CoV-2 Infection. J Immunol 205, 915–922. 10.4049/jimmunol.2000583.

91. Tiller, T., Busse, C.E., and Wardemann, H. (2009). Cloning and expression of murine Ig genes from single B cells. J Immunol Methods 350, 183–193. 10.1016/j.jim.2009.08.009.

92. Ehlers, M., Fukuyama, H., McGaha, T.L., Aderem, A., and Ravetch, J.V. (2006). TLR9/MyD88 signaling is required for class switching to pathogenic IgG2a and 2b autoantibodies in SLE. J Exp Med 203, 553–561. 10.1084/jem.20052438.

93. Ho, I.Y., Bunker, J.J., Erickson, S.A., Neu, K.E., Huang, M., Cortese, M., Pulendran, B., and Wilson, P.C. (2016). Refined protocol for generating monoclonal antibodies from single human and murine B cells. J Immunol Methods 438, 67–70. 10.1016/j.jim.2016.09.001.

94. Davis, C.W., Jackson, K.J.L., McElroy, A.K., Halfmann, P., Huang, J., Chennareddy, C., Piper, A.E., Leung, Y., Albarino, C.G., Crozier, I., et al. (2019). Longitudinal Analysis of the Human B Cell Response to Ebola Virus Infection. Cell 177, 1566–1582 e1517. 10.1016/j.cell.2019.04.036.

95. Punjani, A., Rubinstein, J.L., Fleet, D.J., and Brubaker, M.A. (2017). cryoSPARC: algorithms for rapid unsupervised cryo-EM structure determination. Nature methods 14, 290--296.

96. Bepler, T., Morin, A., Rapp, M., Brasch, J., Shapiro, L., Noble, A.J., and Berger, B. (2019). Positive-unlabeled convolutional neural networks for particle picking in cryo-electron micrographs. Nature methods 16, 1153--1160.

97. Punjani, A., Zhang, H., and Fleet, D.J. (2020). Non-uniform refinement: adaptive regularization improves single-particle cryo-EM reconstruction. Nature methods 17, 1214--1221.

98. Zivanov, J., Nakane, T., and Scheres, S.H.W. (2020). Estimation of high-order aberrations and anisotropic magnification from cryo-EM data sets in RELION-3.1. IUCrJ 7, 253--267.

99. Jumper, J., Evans, R., Pritzel, A., Green, T., Figurnov, M., Ronneberger, O., Tunyasuvunakool, K., Bates, R., Dek, A., Potapenko, A., and, et al. (2021). Highly accurate protein structure prediction with AlphaFold. Nature 596, 583--589.

100. Mirdita, M., Schtze, K., Moriwaki, Y., Heo, L., Ovchinnikov, S., and Steinegger, M. (2022). ColabFold: making protein folding accessible to all. Nature Methods, 1--4.

101. Pettersen, E.F., Goddard, T.D., Huang, C.C., Couch, G.S., Greenblatt, D.M., Meng, E.C., and Ferrin, T.E. (2004). UCSF Chimera—a visualization system for exploratory research and analysis. Journal of computational chemistry 25, 1605--1612.

102. Dunbar, J., Krawczyk, K., Leem, J., Marks, C., Nowak, J., Regep, C., Georges, G., Kelm, S., Popovic, B., and Deane, C.M. (2016). SAbPred: a structure-based antibody prediction server. Nucleic acids research 44, W474--W478.

103. Wang, R.Y.-R., Song, Y., Barad, B.A., Cheng, Y., Fraser, J.S., and DiMaio, F. (2016). Automated structure refinement of macromolecular assemblies from cryo-EM maps using Rosetta. Elife 5, e17219.

104. Chen, V.B., Arendall, W.B., Headd, J.J., Keedy, D.A., Immormino, R.M., Kapral, G.J., Murray, L.W., Richardson, J.S., and Richardson, D.C. (2010). MolProbity: all-atom structure validation for macromolecular crystallography. Acta Crystallographica Section D: Biological Crystallography 66, 12--21.

105. Barad, B.A., Echols, N., Wang, R.Y.-R., Cheng, Y., DiMaio, F., Adams, P.D., and Fraser, J.S. (2015). EMRinger: side chain--directed model and map validation for 3D cryo-electron microscopy. Nature methods 12, 943--946.

106. Emsley, P., and Crispin, M. (2018). Structural analysis of glycoproteins: building N-linked glycans with Coot. Acta Crystallographica Section D: Structural Biology 74, 256--263.

107. Liebschner, D., Afonine, P.V., Baker, M.L., Bunkczi, G., Chen, V.B., Croll, T.I., Hintze, B., Hung, L.W., Jain, S., McCoy, A.J., and, et al. (2019). Macromolecular structure determination using X-rays, neutrons and electrons: recent developments in Phenix. Acta Crystallographica Section D: Structural Biology 75, 861--877.

108. Moriarty, N.W., Grosse-Kunstleve, R.W., and Adams, P.D. (2009). electronic Ligand Builder and Optimization Workbench (eLBOW): a tool for ligand coordinate and restraint generation. Acta Crystallographica Section D: Biological Crystallography 65, 1074--1080.

109. Agirre, J., Iglesias-Fernndez, J., Rovira, C., Davies, G.J., Wilson, K.S., and Cowtan, K.D. (2015). Privateer: sorware for the conformational validation of carbohydrate structures. Nature structural \& molecular biology 22, 833--834.

110. Pettersen, E.F., Goddard, T.D., Huang, C.C., Meng, E.C., Couch, G.S., Croll, T.I., Morris, J.H., and Ferrin, T.E. (2021). UCSF ChimeraX: Structure visualization for researchers, educators, and developers. Protein science : a publication of the Protein Society 30, 70–82. 10.1002/pro.3943.

111. Schlee, S., Straub, K., Schwab, T., Kinateder, T., Merkl, R., and Sterner, R. (2019). Prediction of quaternary structure by analysis of hot spot residues in protein-protein interfaces: the case of anthranilate phosphoribosyltransferases. Proteins 87, 815–825. 10.1002/prot.25744.

112. Bianchi, M., Turner, H.L., Nogal, B., Cottrell, C.A., Oyen, D., Pauthner, M., Bastidas, R., Nedellec, R., McCoy, L.E., Wilson, I.A., and, et al. (2018). Electron-microscopy-based epitope mapping defines specificities of polyclonal antibodies elicited during HIV-1 BG505 envelope trimer immunization. Immunity 49, 288--300.

113. Antanasijevic, A., Sewall, L.M., Cottrell, C.A., Carnathan, D.G., Jimenez, L.E., Ngo, J.T., Silverman, J.B., Groschel, B., Georgeson, E., Bhiman, J., and, et al. (2021). Polyclonal antibody responses to HIV Env immunogens resolved using cryoEM. Nature communications 12, 1--17.

114. Clements, J.D., and Norton, E.B. (2018). The Mucosal Vaccine Adjuvant LT(R192G/L211A) or dmLT. mSphere 3. 10.1128/mSphere.00215-18.

115. Cheng, A., Negro, C., Bruhn, J.F., Rice, W.J., Dallakyan, S., Eng, E.T., Waterman, D.G., Potter, C.S., and Carragher, B. (2021). Leginon: New features and applications. Protein Science 30, 136--150.

116. Kimanius, D., Dong, L., Sharov, G., Nakane, T., and Scheres, S.H.W. (2021). New tools for automated cryo-EM single-particle analysis in RELION-4.0. Biochemical Journal 478, 4169--4185.

117. He, L., Diedrich, J., Chu, Y.-Y., and Yates Iii, J.R. (2015). Extracting accurate precursor information for tandem mass spectra by RawConverter. Analytical chemistry 87, 11361--11367.

118. Xu, T., Park, S.K., Venable, J.D., Wohlschlegel, J.A., Diedrich, J.K., Cociorva, D., Lu, B., Liao, L., Hewel, J., Han, X., et al. (2015). ProLuCID: An improved SEQUEST-like algorithm with enhanced sensitivity and specificity. J Proteomics 129, 16–24. 10.1016/j.jprot.2015.07.001.

119. Tabb, D.L., McDonald, W.H., and Yates, J.R. (2002). DTASelect and Contrast: tools for assembling and comparing protein identifications from shotgun proteomics. Journal of proteome research 1, 21--26.

120. Peng, J., Elias, J.E., Thoreen, C.C., Licklider, L.J., and Gygi, S.P. (2003). Evaluation of multidimensional chromatography coupled with tandem mass spectrometry (LC/LC-MS/MS) for large-scale protein analysis: the yeast proteome. Journal of proteome research 2, 43--50.

121. Baboo, S., Diedrich, J.K., and Martnez, B. (2021). DeGlyPHER: an ultrasensitive method for the analysis of viral spike N-glycoforms. Analytical Chemistry 93, 13651--13657.

122. Mirdita, M., Schutze, K., Moriwaki, Y., Heo, L., Ovchinnikov, S., and Steinegger, M. (2022). ColabFold: making protein folding accessible to all. Nat Methods 19, 679–682. 10.1038/s41592-022-01488-1.

123. Fleckenstein, J.M., and Roy, K. (2009). Purification of recombinant high molecular weight two-partner secretion proteins from Escherichia coli. Nat Protoc 4, 1083–1092. 10.1038/nprot.2009.87.

124. Chakraborty, S., Randall, A., Vickers, T.J., Molina, D., Harro, C.D., DeNearing, B., Brubaker, J., Sack, D.A., Bourgeois, A.L., Felgner, P.L., et al. (2019). Interrogation of a live-attenuated enterotoxigenic Escherichia coli vaccine highlights features unique to wild-type infection. NPJ Vaccines 4, 37. 10.1038/s41541-019-0131-7.

